# Sulfur Amino Acid Restriction Enhances Exercise Capacity in Mice by Boosting Fat Oxidation in Muscle

**DOI:** 10.1101/2024.06.27.601041

**Authors:** Charlotte G Mann, Michael R MacArthur, Jing Zhang, Songlin Gong, Jenna E AbuSalim, Craig J. Hunter, Wenyun Lu, Thomas Agius, Alban Longchamp, Florent Allagnat, Joshua Rabinowitz, James R Mitchell, Katrien De Bock, Sarah J Mitchell

## Abstract

Dietary restriction of the sulfur-containing amino acids methionine and cysteine (SAAR) improves body composition, enhances insulin sensitivity, and extends lifespan; benefits seen also with endurance exercise. Yet, the impact of SAAR on skeletal muscle remains largely unexplored. Here we demonstrate that one week of SAAR in sedentary, young, male mice increases endurance exercise capacity. Indirect calorimetry showed that SAAR increased lipid oxidation at rest and delayed the onset of carbohydrate utilization during exercise. Transcriptomic analysis revealed increased expression of genes involved in fatty acid catabolism especially in glycolytic muscle following SAAR. These findings were functionally supported by increased fatty acid circulatory turnover flux and muscle β-oxidation. Reducing lipid uptake from circulation through endothelial cell (EC)-specific CD36 deletion attenuated the running phenotype. Mechanistically, VEGF-signaling inhibition prevented exercise increases following SAAR, without affecting angiogenesis, implicating noncanonical VEGF signaling and EC CD36-dependent fatty acid transport in regulating exercise capacity by influencing muscle substrate availability.

## Introduction

Caloric restriction (CR) is the gold standard to increase life and health span in various model organisms (Fontana and Partridge, 2015). The reduction of calories was first shown to increase lifespan in mammals early in the 20th century (McCay et al., 1935; Osborne et al., 1917). Ever since, different modalities and degrees of restriction have been introduced and studied in their efficacy in extending health- and lifespan (Lee et al., 2021). Dietary restriction of the sulfur-containing amino acids methionine and cysteine (SAAR) improves body composition, reverses insulin resistance and extends lifespan in rodents (Miller et al., 2005; Orentreich et al., 1993) by eliciting strong metabolic effects. In brown adipose tissue (BAT), SAAR increases thermogenesis and energy expenditure (EE), while also increasing lipolysis and oxidative phosphorylation in white adipose tissue (WAT) and the liver (Forney et al., 2020; Hasek et al., 2010; Patil et al., 2015). The effects of SAAR in skeletal muscle have been less well studied.

Due to its large overall mass and high metabolic activity, skeletal muscle plays crucial roles in the maintenance of systemic homeostasis. Skeletal muscles differ in fiber type composition, traditionally categorized by their myosin heavy-chain content, but also in their metabolic profiles and energy substrate preferences (Egan and Zierath, 2013; Rowe et al., 2014). The extensor digitorum longus (EDL) and the soleus, are characterized as fast and slow twitch muscles respectively, as they use glycolysis versus oxidative phosphorylation as the dominant energy source (Brooke and Kaiser, 1970; Brown, 1973). Muscle fibers adapt to endurance exercise training, enabling better responses to future challenges. These adaptations include the promotion of fiber type transformation (from type IIb/IId/x to IIa), mitochondrial biogenesis, enhanced insulin sensitivity and improved metabolic flexibility. It has been reported that muscle vascularization correlates with mitochondrial density and oxidative capacity (Haas and Nwadozi, 2015) and endurance exercise stimulates angiogenesis in skeletal muscle. Interestingly, the increase in insulin sensitivity, muscle angiogenesis and improved metabolic flexibility are also observed after SAAR, but whether and how SAAR affects skeletal muscle metabolic homeostasis is not known.

Previous research has shown that SAAR increases angiogenesis in skeletal muscle (Longchamp et al., 2018). Angiogenesis is a crucial adaptive response in both developmental and pathophysiological conditions characterized by insufficient oxygen and nutrient supply (Potente et al., 2011). Endurance exercise is one of the few non-pathological settings of vascular expansion during adulthood. During angiogenesis transcription factors like PGC1α/β, estrogen-related receptor (ERR) α/γ, and activating transcription factor 4 (ATF4), are induced by various stimuli including mechano-stress responses or the integrated stress response (ISR). The ISR can be triggered by either endoplasmic reticulum (ER) stress or amino acid (AA) deprivation (Abcouwer et al., 2002; Fan et al., 2021). These adaptive responses play pivotal roles in regulating muscle metabolism and vascular density by enhancing vascular endothelial growth factor (VEGF) levels (Arany et al., 2008; Gorski and Bock, 2019; Matsakas et al., 2012; Narkar et al., 2011; Rowe et al., 2011). VEGF-A is the primary regulator of angiogenesis. By binding to the receptor VEGFR2 VEGF-A initiates a cascade of signal-transduction involving pathways including phosphoinositide 3-kinase (PI3K) and mitogen-activated protein kinase (MAPK), facilitating EC migration, proliferation, and vessel formation (Olsson et al., 2006). VEGF signaling also induces changes in energy metabolism, promoting increased glucose uptake and glycolysis in ECs to meet the energy demands of migration (De Bock et al., 2013). The role of the other VEGF isoforms is less understood. Initially considered to passively regulate angiogenesis by scavenging VEGFR1 (Robciuc et al., 2016), research suggests that VEGF-B may actively modulate EC fatty acid uptake (Dijkstra et al., 2014; Falkevall et al., 2017; Hagberg et al., 2010; Kivelä et al., 2019; Mehlem et al., 2016; Ning et al., 2020).

CR as well as SAAR have been shown to promote revascularization and recovery from femoral artery ligation in rodents (Kondo et al., 2009; Longchamp et al., 2018) and the ability to maintain vascular health in rodents and non-human primates in part by preserving capillary density in skeletal muscle via regulation of VEGF (Omodei and Fontana, 2011). It is unknown whether the SAAR dependent increase in capillary density in skeletal muscle induces functional changes and if this is sufficient to increase exercise performance. Further, the role of VEGF in this process is unexplored.

Here we show that short-term SAAR is sufficient to induce the metabolic benefits of SAAR while also increasing endurance exercise capacity in young, sedentary, male mice. This is achieved by mimicking the metabolic effects of endurance exercise in glycolytic muscle, which requires active lipid transport and occurs through noncanonical VEGF signaling.

## Results

### Short-term SAAR induces shifts in metabolism and increases endurance exercise capacity in young, sedentary, male mice

To evaluate the effect of short-term SAAR on systemic metabolism, we performed SAAR for seven days in young, sedentary, male mice. Metabolic parameters included indirect calorimetry and daily measurement of food intake and body weight. Exercise parameters included a one-time maximal endurance treadmill test on day seven (experimental scheme, Figure 1A). Seven days of SAAR reduced body weight by an average of 8.65% (Figures 1B and S1A) and increased food intake (Figure 1C), consistent with previous reports on SAAR (Miller et al., 2005; Orentreich et al., 1993). Lean to fat mass ratio was not changed by SAAR (Figures S1B-D).

To further understand systemic metabolic changes following seven days of SAAR, we performed indirect calorimetry using metabolic cages, including voluntary running wheels. After seven days, mice on SAAR showed elevated EE during both the active and passive phase (Figures S1E-F). Additionally, seven days of SAAR lowered the respiratory exchange ratio (RER) (Figures 1D-E) without any alterations in voluntary wheel running or overall locomotion (Figures 1F and S1G-H).

To test whether the effects on systemic metabolism translate to functional changes, we measured maximal endurance exercise capacity using a one-time treadmill test to exhaustion. Seven days of dietary SAAR significantly increased endurance exercise capacity in sedentary, young, male mice (Figure 1G), where SAAR mice ran approximately 1.5 times longer than control animals (956.5 ± 306.7 m and 634.8 ± 347.3 m respectively). Running performance was independent of both absolute body weight at testing and relative changes in body weight during studies (Figures S1I-J).

SAAR is reported to have several sexually dimorphic phenotypes (Jonsson et al., 2021; Wanders et al., 2017; Yu et al., 2018), therefore we checked whether the running endurance phenotype showed sexual specificity. Although young female mice did have lower body weight after SAAR (Figure S1K), they did not increase endurance exercise performance (Figure S1L). Due to this sexually dimorphic response, we exclusively used male mice for subsequent studies.

RER calculates substrate utilization during activity (Speakman, 2013) and a lower RER indicates increased reliance on fat oxidation. To directly measure substrate utilization during exercise, we performed a one-time endurance exercise test in metabolic treadmills, monitoring the RER continuously throughout the exercise bout. Consistent with our non-metabolic treadmill data, SAAR mice ran significantly longer (Figures S1M-N). At the terminal spike of the RER as the animals reach exhaustion, the RER was lower, indicating more fat utilization, in SAAR animals compared to controls (Figure 1H).

**Figure 1.**
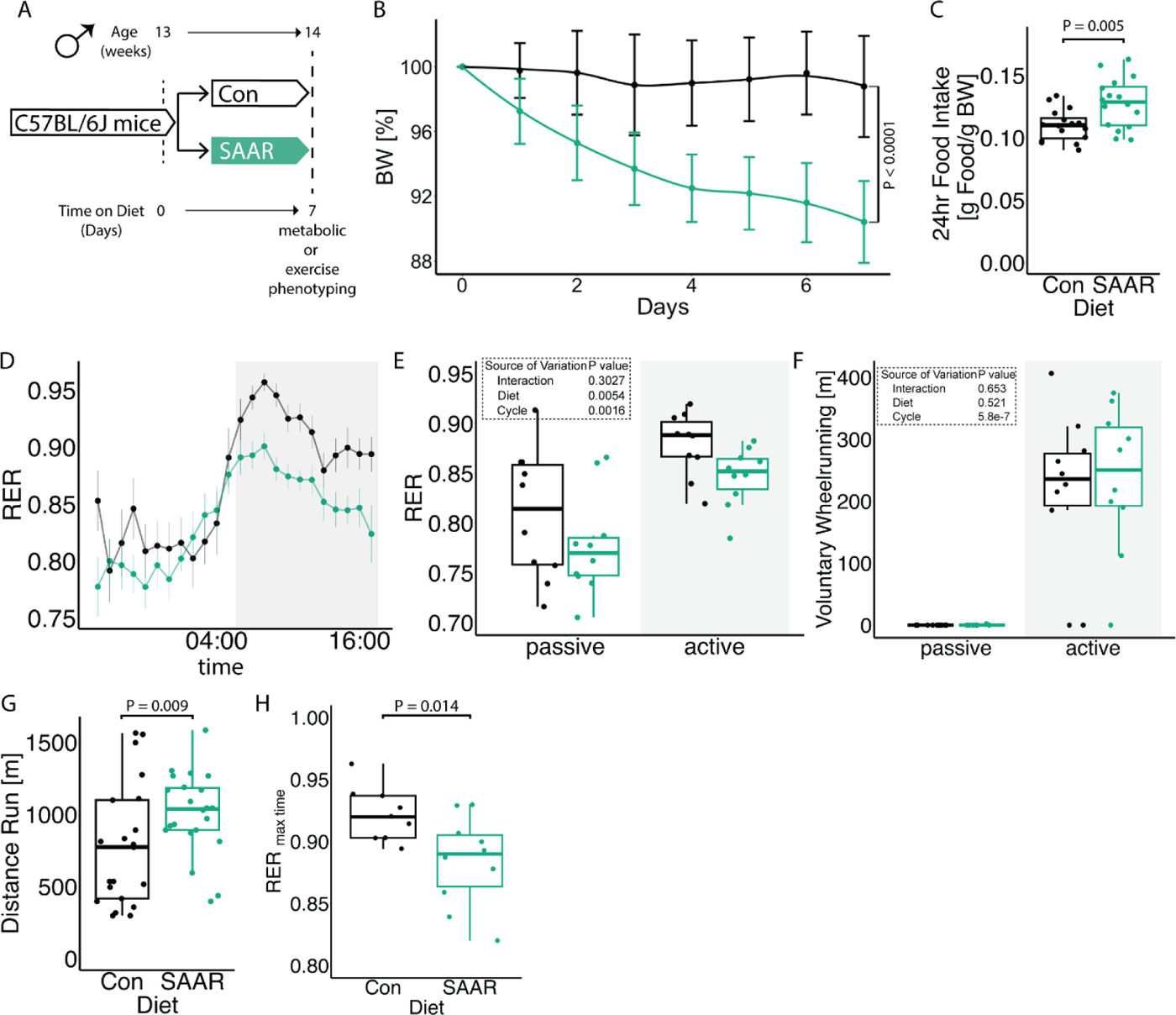
Short-term SAAR induces shifts in metabolism and increases endurance exercise capacity in young, sedentary, male mice. A. Experimental set up and color scheme used throughout figure 1 and figure S1. B. Body weight trajectory over time, shown as percent of starting body weight (n = 10) of male mice given *ad libitum* access to sulfur amino acid restricted (SAAR) versus control (Con) diet for seven days. C. Food intake expressed as grams of food per gram of body weight per mouse within a 24 h period (n = 10) of male mice given *ad libitum* access to SAAR versus Con diet on day seven. D. Sable systems indirect calorimetry measurements of respiratory exchange ratios (CO_2_ emission/O_2_ consumption, VCO_2_/VO_2_, RER) over a 24 h period (n = 10) and E. the average RER during a 12 h – 12 h light–dark cycle (n = 10/group) of male mice given *ad libitum* access to SAAR versus Con diet on day seven. F. The average wheel running in meter during a 12 h – 12 h light–dark cycle (n = 10) of male mice given *ad libitum* access to SAAR versus Con diet on day seven. G. Distance ran in meter during a one-time maximal endurance test (n = 20) of male mice given *ad libitum* access to SAAR versus Con diet on day seven. H. Quantification of RER at maximal exercise time, during a one-time maximal endurance test, performed on a metabolic treadmill (Harvard Apparatus) (n = 10) of male mice given *ad libitum* access to SAAR versus Con diet on day seven. All data is shown as mean and error bars indicate SD unless otherwise noted; p values indicate the significance of the difference by Student’s t test or two-way ANOVA with Sidak’s multiple comparisons test between diets or diet and cycle (indirect calorimetry); significance is determined by a p value of p < 0.05. Each dot represents an individual mouse. See also Figure S1 and Table S1.

### Transcriptomics across muscle depots reveal metabolic shift from glycolytic toward oxidative

Little is known about the effects of SAAR on metabolic capacity in skeletal muscle. Most work has focused on skeletal muscle after long-term SAAR, specifically insulin sensitivity or alterations in skeletal muscle composition with SAAR in aged mice (Ghosh et al., 2017, 2014; Swaminathan et al., 2021). Since our data suggests that seven days of SAAR is sufficient to alter systemic metabolism, we aimed to further characterize metabolic changes in specific skeletal muscles.

To investigate SAAR effects on skeletal muscle while accounting for fiber type, we compared glycolytic EDL with oxidative soleus using bulk RNAseq (experimental scheme, Figure 2A). Gene set overrepresentation analysis on genes affected by diet independent of muscle type showed upregulation of pathways associated with fatty acid or organic acid catabolism and muscle fiber type switching and downregulation of pathways associated with extracellular matrix associated processes and collagen biosynthesis (Figure S2A). The expression of many genes was coordinated in a fiber type dependent fashion (Figures S2B-C), so we investigated expression changes after SAAR within EDL and soleus compared to control diet. Gene-level analysis of fatty acid import and catabolic genes revealed consistently stronger effects in EDL compared to soleus (Figure 2B). A validated ISR/SAAR target gene-set (Torrence et al., 2021) revealed limited muscle type-specific responses, including in the ATF4 target *Cth* (Hine et al., 2015) (Figure S2F). This indicated that depot-specific responses to SAAR are specific for fatty acid metabolism associated genes and may be ISR-independent.

We next looked specifically for expression patterns that showed a diet-by-muscle depot interaction. Significant positive interaction terms (enriched in EDL but not soleus after diet) included the previously identified organic acid catabolic processes and β-oxidation, as well as mitochondrial matrix, (Figure 2C) suggesting a transcriptomic shift of glycolytic EDL to a more oxidative phenotype. TCA cycle enzymes showed similar muscle depot-specific responses on the transcriptomic level (Figure 2D). We also confirmed transcriptomic changes at the protein level, finding that seven days of SAAR was sufficient to increase electron transport chain (ETC) complexes in both EDL and soleus (Figures 2E and S2D-E). The most pronounced changes were observed in SHDB which was increased by approximately 50% after SAAR in EDL (Figure 2F) consistent with changes observed at the transcript level.

**Figure 2.**
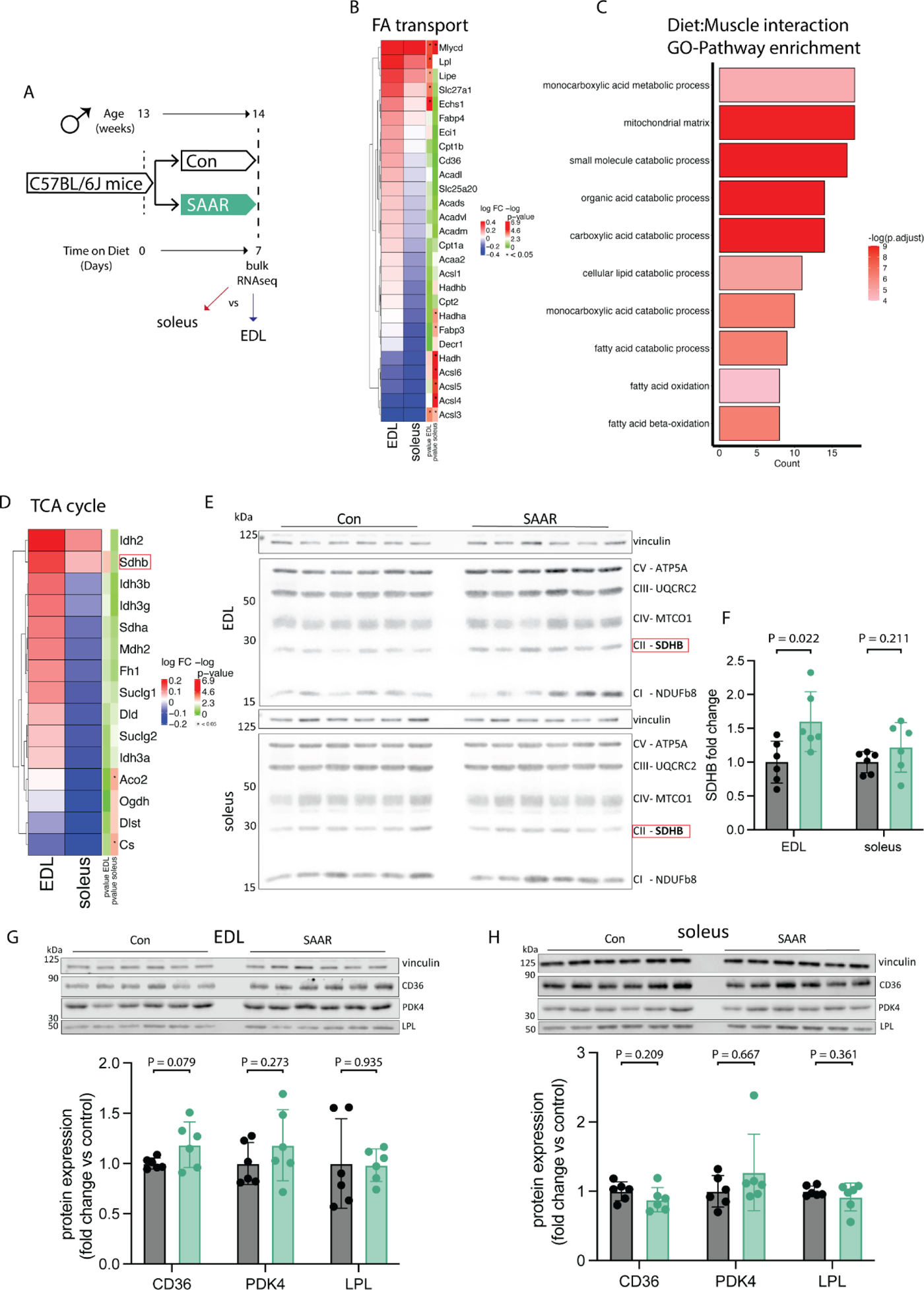
Transcriptomics across muscle depots reveal metabolic shift from glycolytic toward oxidative. A. Experimental set up and color scheme used throughout figure 2 and figure S2. B. Fold changes of transcripts associated with fatty acid (FA) catabolism and transport as identified in supplementary figure 2A in muscle of male mice (n = 6) given *ad libitum* access to sulfur amino acid restricted (SAAR) versus control (Con) diet for seven days. C. Pathway enrichment analysis of genes showing significant diet by muscle interaction effects. D. Fold changes (SAAR vs Con) of TCA cycle genes in EDL and soleus. E. Representative blots of electron transport chain complexes and F. quantification of relative protein abundance normalized to vinculin of SDHB for EDL and soleus (n = 6) of male mice given *ad libitum* access to SAAR versus Con diet on day seven. G. Representative blots of CD36, LPL and PDK4 and vinculin (**top**) and quantification of relative protein abundance normalized to vinculin of CD36, PDK4, and LPL (**bottom**) in EDL (n = 6) of male mice given *ad libitum* access to SAAR versus Con diet on day seven. H. Representative blots of CD36, LPL and PDK4 and vinculin (**top**) and quantification of relative protein abundance normalized to vinculin of CD36, PDK4, and LPL from blots (**bottom**) in soleus (n = 6) of male mice given *ad libitum* access to SAAR versus Con diet on day seven. All data is shown as mean and error bars indicate SD unless otherwise noted; p values indicate the significance of the difference by Student’s t test between diets; significance is determined by a p value of p < 0.05. See also Figure S2 and Table S2.

To determine the overlap between transcriptional adaptation to endurance training and short-term SAAR we compared the transcriptional response to either training or SAAR in gene sets associated with oxidative phosphorylation, using a recently published dataset (Furrer et al., 2023). Transcriptional regulation of both TCA cycle genes and mitochondrial matrix genes showed overlapping patterns between training and EDL diet response, whereas this was not observed to the same extent in the soleus diet response (Figures S2G-H).

We also assessed changes in the protein levels of metabolic enzymes regulating energy homeostasis (CD36, PDK4 and LPL), that were upregulated at the transcriptomic level, using western blot in both the EDL and soleus after seven days of SAAR. CD36 trended towards significant increase after SAAR in EDL only, consistent with the transcriptomic data. LPL and PDK4 showed non-significant increases after SAAR in EDL, however protein levels were variable overall (Figure 2G). Neither CD36, LPL nor PDK4 showed changes after SAAR in soleus (Figure 2H). These changes prompted us to investigate the functional relationship between circulatory lipid turnover and handling.

### SAAR increases muscle lipid flux without altering lipid pool sizes

**Figure 3.**
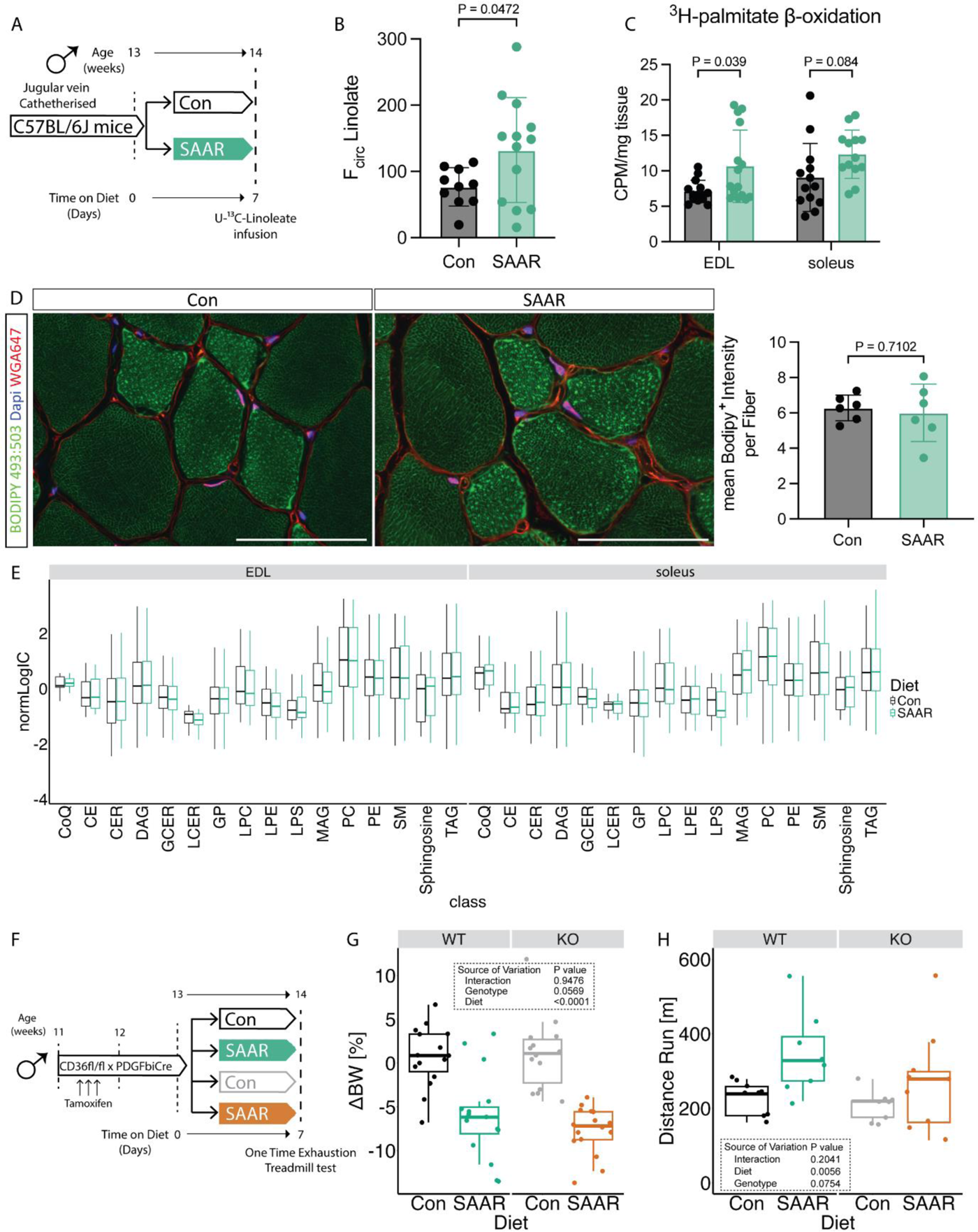
SAAR increases muscle lipid flux without altering lipid pool sizes. A. Experimental design and color scheme used in figure 3A-E and figure S3A. B. Circulatory carbon flux (n = 10–13) of ^13^C_18_-U-Linolate of jugular vein catheterized male mice given *ad libitum* access to sulfur amino acid restricted (SAAR) versus control (Con) diet for seven days. C. *Ex vivo* β-oxidation measured by incorporation of ^3^H-palmitic acid in ^3^H-H_2_O in muscles of male mice fed a Con or SAAR diet for seven days (n = 15). D. Representative fluorescence images (**left**) of BODIPY 493:503 (green), WGA647 (red) and dapi (blue) staining in EDL cross-sections (scale bar, 50 μm) and quantification of Bodipy^+^ Intensity within fibers (**right**) of male mice fed a Con or SAAR for seven days (n = 6). E. Lipidomics analysis from muscle of male mice fed a Con or SAAR diet for seven days (n = 6), summarized as normalized ion counts of each main lipid class. F. Experimental set up and color scheme used in figure 3 G-H and figure S3D-I. G. Percent change in body weight (n = 16) of male WT and EC^CD36-/-^ mice given *ad libitum* access to SAAR versus Con diet after seven days. H. Distance ran during a one-time maximal endurance test (n = 8) of male WT and EC^CD36-/-^ mice given *ad libitum* access to SAAR versus Con diet on day seven. All data is shown as mean and error bars indicate SD unless otherwise noted; p values indicate the significance of the difference by Student’s t test between diets, or two-way ANOVA with Sidak’s multiple comparisons test between diets and muscle or genotype; significance is determined by a p value of p < 0.05. See also Figure S3 and Table S3-5.

To test whether the transcript and protein level changes we observed (Figures 2B-H) resulted in functional alterations in lipid metabolism, we measured the circulatory turnover flux (F_circ_) of linoleate in steady-state infusions (Hui et al., 2020) in jugular vein-catheterized mice after seven days of SAAR (experimental scheme, Figure 3A). Linoleate turnover flux was significantly increased after SAAR compared to control mice (Figure 3B). To test whether skeletal muscle fatty acid consumption is involved in driving increased flux, we assessed *ex vivo* skeletal muscle β-oxidation after short-term SAAR, using radiolabeled palmitate in EDL and soleus. Across both muscle depots, there was a significant main effect of diet on β-oxidation, however post-hoc tests revealed a significant control vs SAAR difference in EDL only, highlighting the more pronounced shift of glycolytic muscle to increase lipid oxidation (Figure 3C).

Increased lipid turnover and oxidation can be driven by two major mechanisms: increased circulating lipid concentrations driving an increase in uptake and storage by mass action (Li et al., 2022) or changes in lipid transporter activity. To test if mass action was driving increased turnover and β-oxidation in the muscle, we first measured intramyocellular lipid storage after seven days of diet. Bodipy staining in sections of EDL did not show increased intramyocellular lipid storage between diet groups (Figure 3D). We also performed lipidomic and metabolomic measurements to investigate changes in pool sizes of lipid classes and free fatty acid species after SAAR in tissues. No major lipid classes in either EDL or soleus were affected by seven days of SAAR compared to control (Figure 3E). When assessing differences driven by diet, tissue, or diet and tissue interaction at an individual lipid species level, the dominant source of variance was tissue. However, when focusing on the main effect of diet or diet-by-muscle depot interaction effects, only a small number (3 and 16 respectively) out of 690 measured lipid species were significantly changed (Supplemental Table 3). This trend was also generally true for polar metabolites (Supplemental Table 4). Many metabolites showed significant main effects of muscle depot, for example carnosine and anserine were enriched in glycolytic tissue as previously reported (Luo et al., 2023) (Supplemental Table 4). We also observed a main effect of diet in multiple metabolites associated with dietary protein restriction (ophthalmic acid) (MacArthur et al., 2022) or *Pparg* associated changes in thermogenesis (aminoisobutyric acid) (Roberts et al., 2014), however none of these changes showed diet by muscle depot interaction effects. Downstream metabolites of the transsulfuration pathway (for example taurine) were depleted in the muscle of mice on SAAR (Supplemental Table 4) suggesting SAAR also affects metabolites downstream of methionine in the muscle. Overall, we did not observe any changes due to diet in the pool-size of various free fatty acid species measured in our metabolomic dataset (Figure S3A, supplemental table 4), suggesting that the increased lipid turnover flux is not driven by an increase in lipid availability.

Movement of circulating fatty acids to the skeletal muscle requires their transfer across the EC barrier and EC CD36 facilitates tissue fatty acid uptake (Peche et al., 2023; Son et al., 2018). Since *Cd36* was upregulated by SAAR specifically in the EDL on both the transcriptomic and protein level (Figures 2B,G-H), we tested a potential requirement for increased lipid shuttling into the muscle via EC CD36. To explore this hypothesis, we generated a tamoxifen-inducible EC specific CD36 knockout by crossing mice with a *Cd36* floxed allele with mice hemizygously expressing PDGFβiCre (EC^CD36-/-^) (Claxton et al., 2008). Cre^+^ and Cre^-^ litter mates were distributed equally between control and SAAR group (experimental scheme, Figure 3F). One cohort was used to study metabolic phenotypes and another was used to perform a one-time endurance exercise test. The KO efficiency was confirmed by flow cytometry, gating for CD31^+^CD36^+^ populations in both skeletal muscle and BAT (Figures S3B-E). EC *Cd36* KO did not affect the body weight changes induced by SAAR (Figures 3G and S3F). When performing the endurance exercise capacity test, SAAR significantly increased running performance in WT animals only. Post hoc testing showed increased running performance after SAAR in WT animals and EC^CD36-/-^ deletion attenuated the running performance increases after SAAR (Figure 3H). To test whether EC^CD36-/-^ had an effect on fatty acid composition in skeletal muscle and circulation after one week of diet, we performed bulk metabolomics. After false-discovery rate (FDR) correction, very few metabolites were significantly affected by genotype or diet in EDL (13 and 7 respectively), soleus (3 and 3 respectively) or serum (16 and 28 respectively), none of which showed consistency, likely due to high sample variability (Figures S3G-H, supplementary table 5).

### FGF21 is dispensable for increased running capacity upon SAAR

FGF21 plays a critical role in the systemic metabolic adaptations to SAAR and can drive increased β-oxidation, particularly in adipose tissue (Agius et al., 2024; Forney et al., 2020; Hill et al., 2019; Wanders et al., 2017). To test whether FGF21 is required for increased running performance upon SAAR, mice with an *Fgf21* floxed allele were crossed with CMV-Cre to achieve stable whole body *Fgf21* knock out (FGF21KO) (experimental scheme, Figure 4A). FGF21KO and WT littermates were evenly distributed between dietary groups with one cohort used to assess metabolic changes and a second cohort for one time treadmill testing after seven days of diet. No significant effects of genotype on SAAR-induced body weight changes were observed (Figures 4B and S4A). However, in agreement with previous observations, the decreased body weight in FGF21KO animals on SAAR can be mainly explained by a decrease in food intake (Figure S4B) (Wanders et al., 2017). FGF21KO had no effect on running performance, with both genotypes tending to increase after SAAR (Figure 4C). However, variability in the control groups limited these trends from reaching statistical significance in either genotype (Figure 4C). KO efficiency was confirmed by serum FGF21 ELISA, where SAAR increased circulating FGF21 levels, and no FGF21 was detected in serum of FGF21KO animals (Figure S4C).

We and others have observed increases in FGF21 after SAAR, therefore we tested whether FGF21 alone is sufficient to increase running performance by infusing recombinant FGF21 for seven days. We implanted WT C57BL/6J male mice with osmotic minipumps loaded with either saline solution or recombinant FGF21 (dosed at 1 mg/kg/day) for seven days (experimental scheme, Figure 4D). Expression of known FGF21 downstream targets in BAT were tested using qPCR to confirm the sufficiency of exogenous FGF21 to induce molecular changes. *Ucp1* and *Fgf21* were upregulated in both the exogenously supplied FGF21 group as well as the SAAR group (Figure S4E). Exogenous FGF21 caused a significant reduction in body weight, although to a lesser extent than what we observe with the SAAR diet (Figures 4E and S4D). On day seven of the intervention the animals underwent a one-time treadmill test. Exogenous FGF21 supply was not sufficient to mimic SAAR in increasing endurance exercise capacity (Figure 4F). Taken together FGF21 is not necessary for the increased endurance exercise capacity after SAAR and is not sufficient to increase endurance exercise capacity on its own.

**Figure 4.**
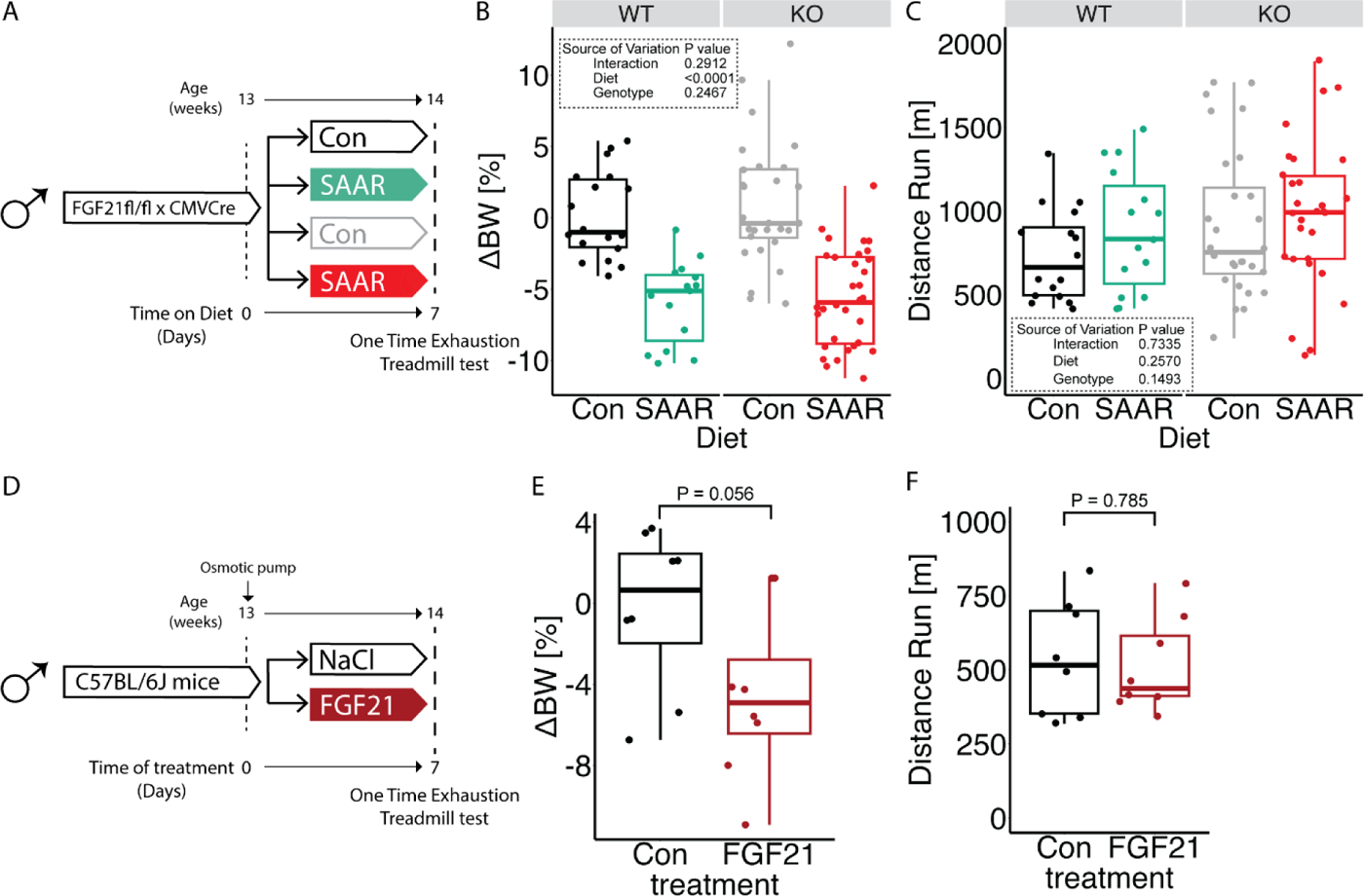
FGF21 is dispensable for increased running capacity upon SAAR. A. Experimental design and color scheme used in figure 4B-C and figure S4A-C. B. Percent change in body weight (n = 11 - 24) of male WT or FGF21KO mice given *ad libitum* access sulfur amino acid restricted (SAAR) versus control (Con) diet for seven days. C. Distance ran during a one-time maximal endurance test (n = 11 - 24) of male WT or FGF21KO mice given *ad libitum* access to SAAR versus Con diet on day seven. D. Experimental set up and color scheme used throughout E-F and figure S4D-E. E. Percent change in body weight (n = 8) of NaCl or recombinant FGF21 treated male mice for seven days. F. Distance ran during a one-time maximal endurance test (n = 8) of NaCl or recombinant FGF21 treated male mice on day seven. All data is shown as mean and error bars indicate SD unless otherwise noted; p values indicate the significance of the difference by Student’s t test between treatments, or two-way ANOVA with Sidak’s multiple comparisons test between diets and genotype; significance is determined by a p value of p < 0.05. See also Figure S4.

### Inhibition of VEGFR signaling blocks increased endurance exercise capacity by SAAR

Our previous work demonstrated that long-term SAAR increases EC VEGF levels and induces angiogenesis in the muscle (Longchamp et al., 2018), which is crucial for exercise performance (Delavar et al., 2014; Olfert et al., 2009). Therefore, we examined the transcriptomic signature associated with VEGF-signaling in our RNA seq dataset. Interestingly we observed that *Vegfb* was increased by SAAR specifically in the EDL (Figures 5A-B) and *Flt1* (encoding VEGFR1) trended towards an increase in both EDL and soleus (Figures 5A,C). *Vegfa* and *Kdr* (encoding VEGFR2) were not affected at the transcript level by short-term SAAR (Figures S5A-B). To test whether induction of VEGF-dependent angiogenesis was a driver of increased running capacity, we treated animals with the pan-VEGFR inhibitor axitinib or vehicle control by oral gavage for the seven-day duration of the dietary intervention (experimental scheme, Figure 5D). Treatment with axitinib did not affect the body weight response to SAAR (Figures 5E and S5C). However, upon one-time endurance exercise testing at day seven, axitinib blunted the increased running performance of SAAR mice (Figure 5F). To test whether this was due to the prevention of SAAR-induced angiogenesis we measured vascular density in the muscle. Immunofluorescent labeling of the vasculature in cryosections of EDL did not show any significant changes in vascular area as a function of either SAAR or axitinib treatment (Figures 5G and S5D). Labeling proliferating ECs using EdU also did not show an effect of SAAR on EC proliferation in either EDL or soleus after seven days (Figure S5E). Similarly, quantifying total CD31^+^ cells using FACS analysis, revealed no increase in EC number after seven days of SAAR in either muscle or BAT (Figures S5F-G).

Since angiogenesis was not changed with axitinib treatment or short-term SAAR, we next assessed whether transcriptional regulation of the VEGF/VEGFR genes were altered using RNA sequencing of the EDL from mice treated with axitinib during seven days of SAAR. After SAAR only in combined with axitinib treatment, *Vegfa* and *b* were increased, while the induction of both receptors after short-term SAAR in combination with axitinib was prevented (Figures 5H-I and S5H-I). Interestingly, we also observed that inhibition of VEGF-signaling downregulated fatty acid transport associated transcripts (Figure S5J) but on the other hand further promoted dietary increases in ETC associated genes (Figure S5K), suggesting that VEGF signaling may control fatty acid availability but not oxidation capacity. To differentiate between the different VEGFRs and their contribution to the endurance exercise capacity increase we observed, we treated mice with DC101, a VEGFR2 specific antibody (Arulanandam et al., 2015) or an IgG control and measured metabolic as well as exercise parameters (experimental scheme, Figure 5J). Inhibiting VEGFR2 specifically with DC101, did not alter the dietary effect of SAAR on either reducing body weight or increasing food intake (Figures 5K and S5L-M). However, inhibiting VEGFR2 was sufficient to prevent the SAAR-induced increase in endurance exercise capacity (Figure 5L). Taken together these data implicate a role for VEGF-signaling in modulating endurance exercise capacity after short-term SAAR.

**Figure 5.**
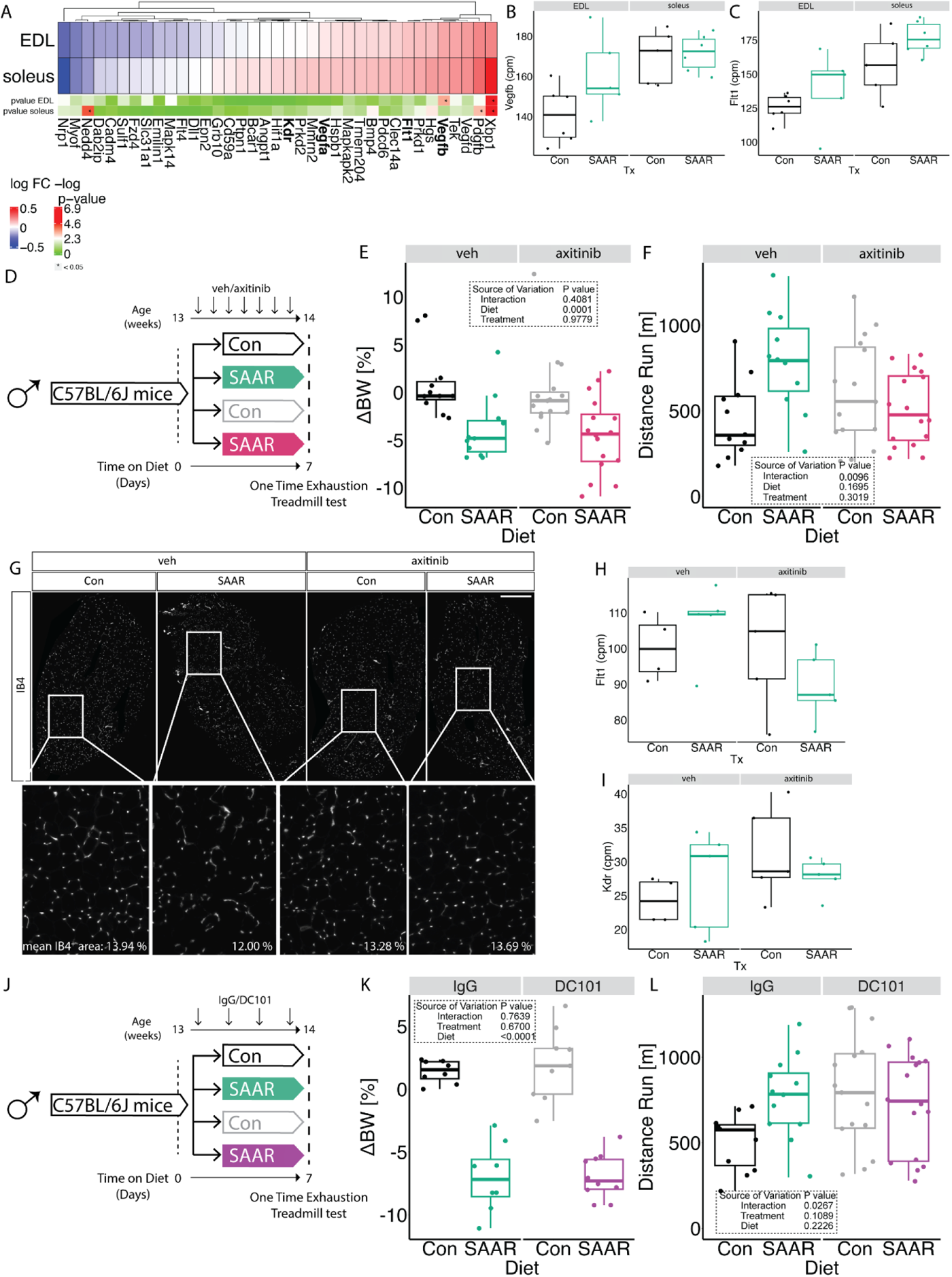
Inhibition of VEGFR signaling blocks increased endurance exercise capacity by SAAR. A. Fold changes of transcripts associated with Vegf signaling using transcriptomic dataset presented in figure 2 in muscles of male mice (n = 6) after sulfur amino acid restriction (SAAR) compared to control (Con) diet for seven days. B. Normalized count values of Vegfb in EDL and soleus from bulkRNA sequencing (n = 6) of male mice given *ad libitum* access to SAAR versus Con diet on day seven. C. Normalized count values of Flt1 in EDL and soleus from bulkRNA sequencing (n = 6) of male mice given *ad libitum* access to SAAR versus Con diet on day seven. D. Experimental set up and color scheme used in figure 5E-F and figure S5C-D. E. Percent change in body weight (n = 16 - 24) of male mice treated with vehicle (veh) or axitinib via oral gavage in combination with *ad libitum* access to SAAR versus Con diet after seven days. F. Distance ran during a one-time maximal endurance test (n = 16-24) of male mice treated with veh or axitinib via oral gavage in combination with *ad libitum* access to SAAR versus Con diet for seven days. G. Representative fluorescence images of IB4 (white) staining in EDL cross-sections of mice fed a Con or SAAR Diet, co-treated with veh or axitinib via oral gavage (scale bar, 400 μm) for seven days (n = 5-8). H. Normalized count values of Vegfb in EDL treated with either veh or axitinib from bulkRNA sequencing (n = 5) of male mice given *ad libitum* access to SAAR versus Con diet on day seven. I. Normalized count values of Flt1 in EDL treated with either veh or axitinib from bulkRNA sequencing (n = 5) of male mice given *ad libitum* access to SAAR versus Con diet on day seven. J. Experimental set up and color scheme used in figure 5K - L and figure S5J - K. K. Percent change in body weight (n = 8-10) of male mice treated with IgG or DC101 via i.p. injection every other day in combination with *ad libitum* access to SAAR versus Con diet after seven days. L. Distance ran during a one-time maximal endurance test (n = 12-16) of male mice treated with IgG or DC101 via i.p injection every other day in combination with *ad libitum* access to SAAR versus Con diet on day seven. All data is shown as mean and error bars indicate SD unless otherwise noted; p values indicate the significance of the difference by two-way ANOVA with Sidak’s multiple comparisons test between diets and treatment; significance is determined by a p value of p < 0.05. See also Figure S5 and Table S6.

## Discussion

Using healthy, sedentary, male mice, we report that short-term sulfur amino acid restriction rewires systemic and muscle metabolism and increases endurance exercise capacity independently of angiogenesis, through noncanonical VEGF-signaling.

Little research has focused on the skeletal muscle functional outcome of the metabolic changes after SAAR. In aged mice, 18 weeks of SAAR rescued lean mass losses upon age but also prevented muscle hypertrophy after overload (Swaminathan et al., 2021). While running performance or other indicators of muscle fitness were not assessed in that study, the authors did show elevated levels of succinate dehydrogenase activity (SDH) suggesting increased oxidative capacity after SAAR in skeletal muscle. Similar findings on citrate synthase activity and mitochondrial biogenesis have also been observed (Perrone et al., 2012). Functionally, it is widely established that SAAR increases EE (Forney et al., 2020; Hasek et al., 2010; Perrone et al., 2010; Wanders et al., 2017; Yu et al., 2018), however little research mentions whole body substrate utilization (RER) or physical activity. Increases in total activity were measured after 8 weeks of SAAR feeding, where studies (Lees et al., 2014, Yu et al., 2018) reported lowered RER after 5 weeks of a western diet restricted in sulfur amino acids but these data could be influenced by different lipid compositions of the diet.

After seven days, we found that SAAR promoted a metabolic shift towards whole body fat oxidation during rest as well as exercise measured by RER which was reflected by increased expression of genes related to fat oxidation and oxidative phosphorylation in skeletal muscle. Functionally, this increased whole body linolate Fcirc and β-oxidation in skeletal muscle. The activation of fat metabolism was more pronounced in muscle with a higher proportion of glycolytic fibers, such as the EDL. However, we did observe increases in oxidative phosphorylation genes in more oxidative muscles as well, and pathways associated with increases in fatty acid catabolism and organic acid import were increased upon diet rather than being specific for diet:muscle interaction. We thus hypothesize that due to the already high fat oxidative capacity of oxidative muscles, changes in EDL are more prone to elicit relevant changes on a functional level. Overall SAAR leads to an increase in oxidative capacity and expression of genes involved in fatty acid catabolism and oxidation in all muscles.

Seven days of SAAR was sufficient to increase both circulatory fatty acid turn-over as well as muscle specific β-oxidation. We did not see changes in either intramyocellular lipid storage or changes in lipid composition or free fatty acid pool size upon the dietary treatment. This suggests that the increase in fat oxidation was fueled by increased fatty acid uptake from the circulation which requires transendothelial transport. Consistently, the expression and protein content of the fatty acid transporter CD36 was increased upon SAAR. Restricting the acute supply of fatty acid influx via endothelial CD36 deletion during exercise was enough to attenuate the endurance exercise capacity phenotype. This underscores the importance of fatty acid supply, in addition to oxidation, for SAAR to elicit its beneficial effects on endurance exercise capacity. Of note, our observations that preventing transendothelial lipid transport ablated the running phenotype demonstrates necessity, but not sufficiency of CD36 for increasing running performance. However, increased lipid uptake upon SAAR is required to fuel oxidative phosphorylation, and whether or not inhibition of oxidative phosphorylation also attenuates the endurance exercise capacity is yet to be tested.

Previously we have shown that long-term SAAR drives angiogenesis in skeletal muscle via activating VEGF-signaling in ECs, leading to increased vascular density (Longchamp et al., 2018). In this study, one week was not sufficient to stimulate neovascularization in the muscle and is consistent with the timeline for neovascularization following exercise training, which requires 2-3 weeks of consistent training for an observable increase in vascular density (Bloor, 2005). We also did not observe an increase in endothelial cell proliferation. We therefore propose that the improved endurance exercise capacity upon SAAR is driven by a metabolic shift favoring fatty acid transport by ECs and oxidation by skeletal muscle rather than by increased muscle vascularization. Interestingly, inhibition of pan VEGFRs or VEGFR2 alone also inhibited the endurance exercise capacity increase upon SAAR implicating a role of VEGF-signaling on SAAR’s effect on endurance exercise capacity independent of its canonical role in angiogenesis. Our transcriptomic dataset after SAAR and axitinib treatment suggests that the effect of inhibiting VEGF-signaling mainly acts on fatty acid uptake as ETC associated genes and genes involved in fatty acid oxidation remained upregulated. Angiogenesis driven by longer-term SAAR treatment could possibly augment the increased endurance exercise performance further and could be tested in future studies.

Previous studies have shown that VEGF-dependent activation of AKT and AMPK is required for EC proliferation (Nagata et al., 2003; Reihill et al., 2011; Stahmann et al., 2010). AMPK has also been identified as a potential trigger for CD36 translocation (Han et al., 2019) and AMPK is known to induce and upregulate fatty acid oxidation via oxidative phosphorylation in the muscle (Han et al., 2019; Salminen et al., 2017). SAAR studies have shown that SAAR activates AMPK via upregulation of H_2_S in a VEGF-dependent fashion in the endothelium (Longchamp et al., 2018), and more recent work established a role for an H_2_S - AMPK axis in chicken muscle (Li et al., 2023). Induction of H_2_S by dietary restriction is widely established (Hine et al., 2015; Jonsson et al., 2021; Zivanovic et al., 2019), and we too observed transcriptional upregulation of *Cth* - a major enzymatic contributor to endogenous H_2_S production in muscle. Even though we did not further investigate AMPKs role in increasing endurance exercise after short-term SAAR, AMPK could be a potential signaling hub integrating upstream VEGF-signaling with downstream upregulation of oxidative phosphorylation and increased fatty acid uptake. A potential signaling cascade could include EC VEGF - ATF4 - H_2_S - AMPK - CD36 signaling resulting in increased fatty acid import into the muscle via the endothelium, allowing for increased systemic turnover, and at the same time promoting β-oxidation in the muscle thereby increasing endurance capacity in sedentary male mice. Previous studies have focused on EC intrinsic VEGF-signaling, showing upregulation of the VEGF - ATF4 - H_2_S - AMPK in ECs but not in the whole muscle tissue. Based on our data, we cannot exclude VEGF-driven metabolic crosstalk between EC and myofibers upon SAAR.

### Limitation of this study

Due to technical limitations, we were unable to unravel the specific contribution of VEGF-A or -B as the molecular driver of endurance exercise capacity increases after short-term SAAR. We found that both VEGF-A as well as VEGF-B are transcriptionally upregulated after dietary SAAR. Inhibition of VEGFR2, which exclusively binds VEGF-A, prevented increased running performance, suggesting a causal role for VEGF-A and its receptor. Skeletal muscle VEGF has been reported to be crucial for exercise training, as deletion of VEGF blunted exercise capacity in mice (Delavar et al., 2014). We and others have previously shown that SAAR increases endothelial VEGF signaling (Das et al., 2018; Longchamp et al., 2018). However, an increase in VEGF-B could still push increased VEGF-A/VEGFR2 interaction due to competition for VEGFR1 binding. Further research and genetic models will be required to completely address this question.

## Acknowledgements/Funding

This work was supported by the National Institute on Aging (P01AG055369 to SJM), an ETH Zurich Doc.Mobility Fellowship (CGM) and ETH Zurich core funding. We dedicate this work to our late mentor and friend, James R Mitchell. We want to thank the center of PhenoGenomics at EPFL for helping us perform the metabolic running experiments.

## Author Contributions

Conceptualization, C.G.M., M.R.M., K.D.B., J.R.M, and S.J.M.; Methodology, C.G.M., M.R.M., J.Z., S.G., J.E.A., W.L., C.H., T.A.; Resources, A.L., F.A., J.R., K.D.B and S.J.M; Writing, C.G.M., M.R.M., K.D.B., and S.J.M..; Funding Acquisition, C.G.M, M.R.M and S.J.M.

## STAR Methods

### RESOURCE AVAILABILITY

#### Lead Contact

Further information and requests for resources and reagents should be directed to and will be fulfilled by the Lead Contact, Sarah Mitchell.

#### Materials Availability

Reagents used to conduct the research detailed in this manuscript are available on request from the Lead Contact, Sarah Mitchell.

#### Data and Code availability

The authors declare that all the data supporting the findings of this study are available within the article and its Supplementary Information Files.

### KEY RESOUCES TABLE

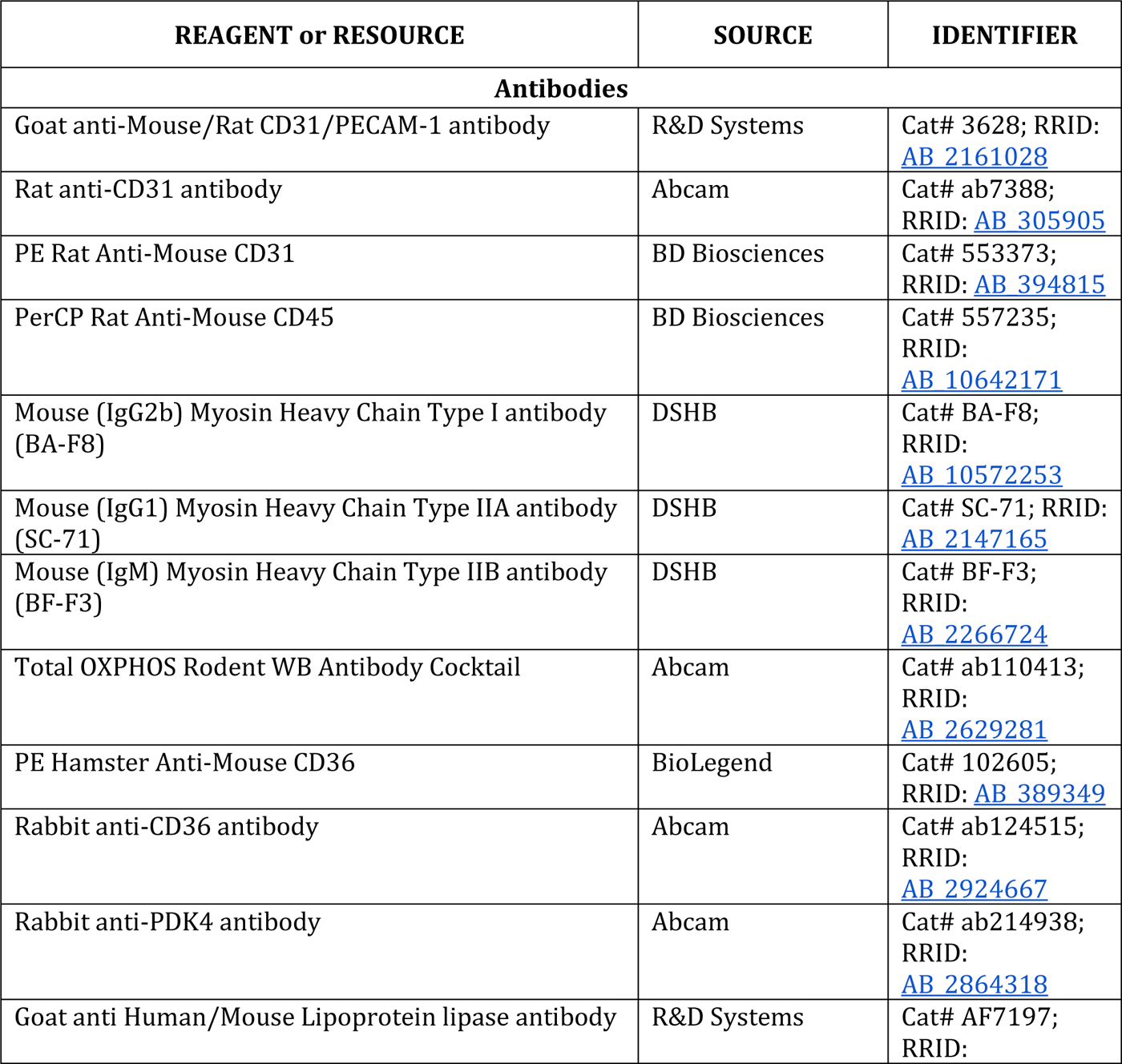
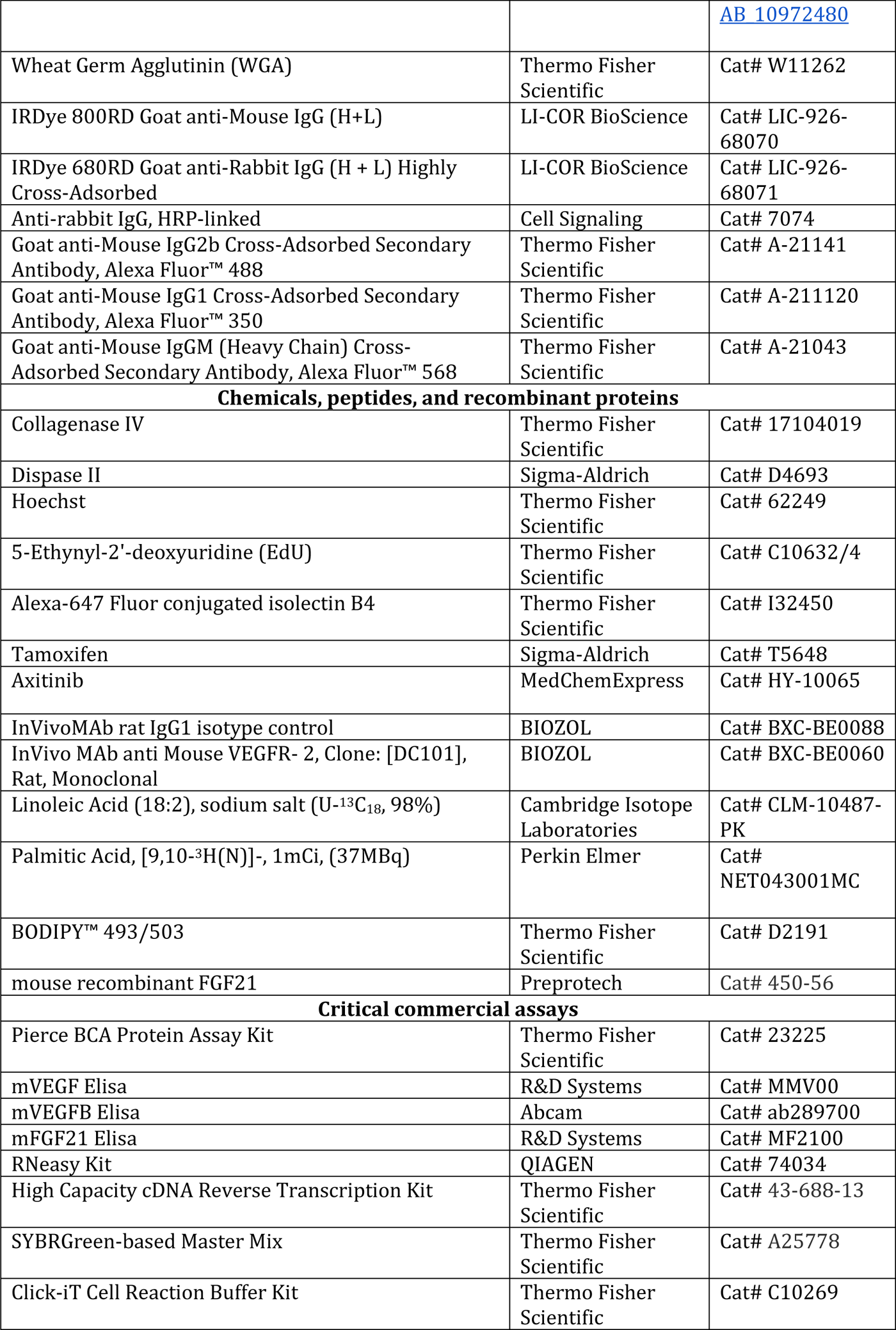
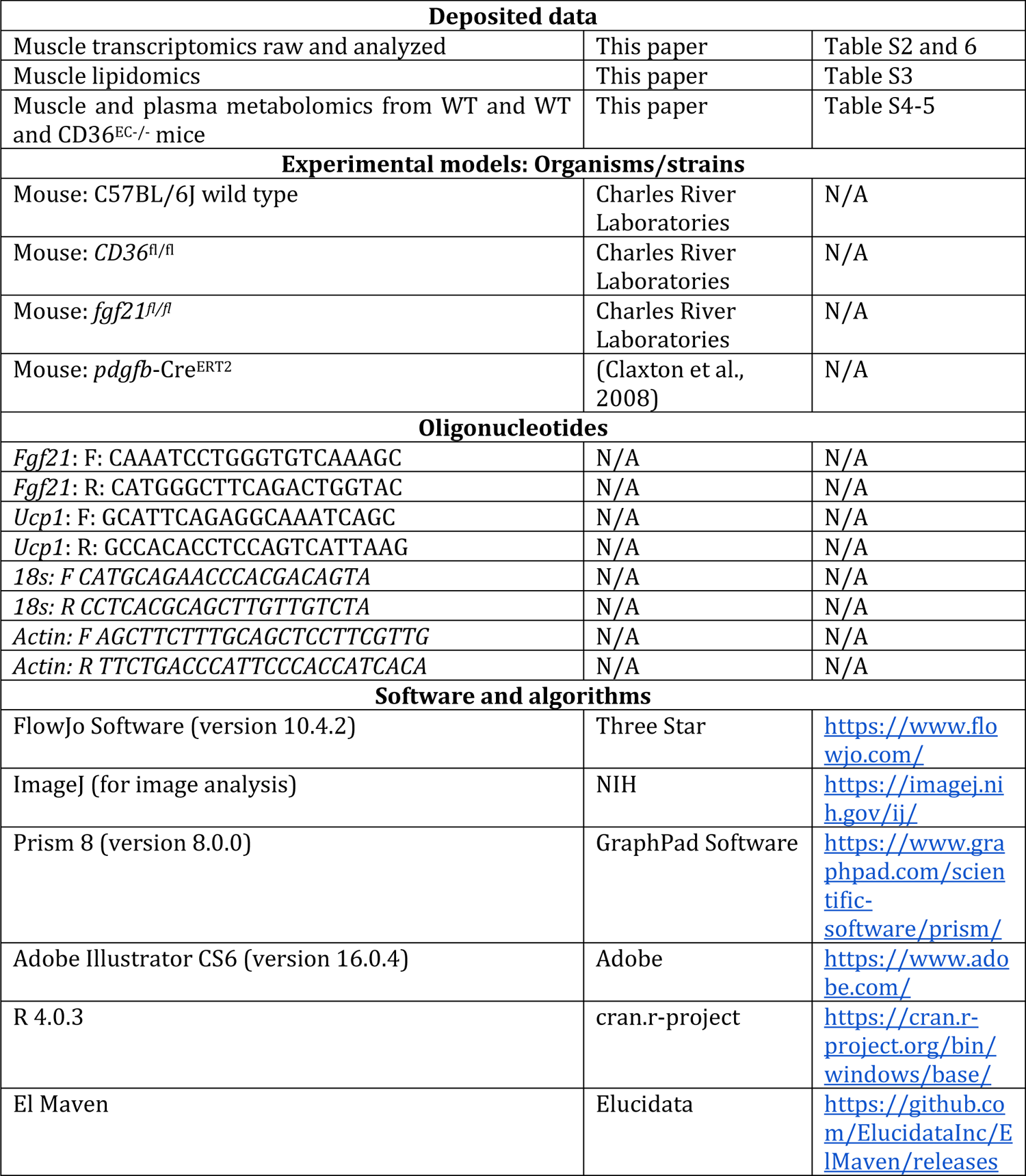

## EXPERIMENTAL MODELS

### EXPERIMENTAL MODEL AND STUDY DETAILS

#### Mice

All animal experiments were approved by the local animal ethics committees (Kantonales Veterinäramt Zürich, licenses ZH211/19, ZH149/21, ZH133/23, Animal Care and Use Committee for Princeton University, Harvard Medical Area or Boston University Institutional Animal Care and Use Committee (IACUC) and Service de la Consommation et des Affaires Vétérinaires SCAV-EXPANIM, license VD-3664), and performed according to local guidelines (TschV, Zurich) and the Swiss animal protection law (TschG). Health status of all mouse lines was regularly monitored according to FELASA guidelines. Mice used in experiments were 8 to 14 weeks old. Mice were housed in standard housing conditions (22 C, 12 h light/dark cycle), with ad *libitum* (AL) access to chow diet (18 % protein, 4.5 % fat, #3437, Provimi Kliba SA) and water.

Wild type (WT) C57BL/6J and *Cd36* LoxP/LoxP mice (*Cd36^tm1.1Ijg^*/J) were purchased from Charles River (Freiburg im Breisgau, Germany). To obtain inducible endothelial cell-specific Cd36 knockout (EC^CD36-/-^) mice, *Cd36* LoxP/LoxP mice were crossed with *PDGFβ.iCreER* mice, an EC-selective inducible Cre-driver line (Claxton et al., 2008). Recombination was induced in 8-10 weeks old male mice by daily intraperitoneal (i.p.) administration of 1mg tamoxifen (T5648, Sigma-Aldrich) dissolved in 1:10 ethanol:corn oil solution for 3 consecutive days. A wash out period of at least seven days was allowed before starting the experiments. Tamoxifen-treated Cre-negative littermates were used as control for all experiments. *Fgf21* knockout (*Fgf21*KO) mice were generated by crossing *Fgf21*LoxP/LoxP mice (B6.129S6(SJL)-Fgf21tm1.2Djm/J) with loxP sites flanking exons 1-3 of the Fgf21 gene with CMV-Cre expressing mice (B6.C-Tg(CMV-Cre)1Cgn/J). The resulting offspring had a deletion in exons 1-3 of Fgf21 in all tissues. The line was subsequently maintained by breeding animals heterozygous for the deletion allele.

Mouse recombinant FGF21 (Cat# 450-56, Peprotech) was dissolved and diluted in sterile distilled water to a final dosage of 1 mg/kg/day. The filled 1007D Alzet osmotic minipump was pre-soaked for 24 hours in NaCl at 37 °C in a dry incubator. Mice were anesthetized with 3% isoflurane in 2 L O2 and kept at 37 °C with an electrical heating pad. A 1-cm incision was made in the skin of the upper back/neck to implant the sterile, preloaded minipump. 5-0 Prolene surgical suture was used to close the wound. Mice received paracetamol (2mg/ml Dafalgan, UPSA) in the drinking water for 48 hours postoperatively.

Aseptic surgery was performed to place catheters in the right jugular vein connected to a vascular access button implanted under the skin on the back of the mouse (Instech Laboratories). Mice were allowed to recover from catheterization surgery for at least 5 days before experimentation. Mice with catheters were individually housed in environmentally enriched cages with AL access to water and food. Catheters were flushed with sterile saline and refilled with sterile heparin glycerol locking solution (SAI Infusion Technologies, HGS) every 5-6 days.

Experimental diets were based on Research Diets D12450J with approximately 18% of calories from protein, 10% from fat and 72% from carbohydrates. SAAR diets containing 1.15g methionine (M)/kg food and lacking cysteine (C) (Miller et al., 2005) in the context of a 17% protein/ 73% carbohydrate calorie diet were provided AL. Food intake was monitored daily during experiments. The Research Diets product number for the control diet is A17101101 and for SAAR diet is A17101103.

Where indicated, axitinib was delivered via daily oral gavage at a dose of 25 mg/kg in 0.5% carboxymethylcellulose vehicle. To block VEGF/VEGFR2 signaling, mice were treated with DC101, a rat monoclonal IgG1 antibody against VEGFR2 (30 μg/kg, i.p., BioXcell) every other day for the duration of the dietary intervention.

To label proliferating cells, an i.p. injection of 5-ethynyl-2’-deoxyuridine (EdU) (E10187, Thermo Fisher Scientific) solution (5 mg/ml in saline, 10 ul/g BW injected) was performed seven hours before sacrificing the mice.

### METHOD DETAILS

#### Exercise experiments

For endurance exercise capacity testing, mice were acclimated to a treadmill system for 3 sessions (5-lane treadmill, Harvard Apparatus, Panlab) before exercise capacity testing. During acclimation sessions each animal ran for 10 min, increasing the speed from 5 m/min up to 10 m/min at minute 5 and kept constant at 10 m/min for 5 min. Thereafter the animal rested for 5 min, followed by 10 min at 10m/min with 5% incline. Following acclimation, mice underwent an aerobic exercise capacity test to exhaustion on a treadmill with 5 degree angle. Mice were motivated to run with a shock grid set at 0.2 mA. Starting speed was 5 m/min for 5 min and was increased by 1 m/min until exhaustion. Speed was capped at a maximum of 20 m/min until exhaustion. Current was taken off the grid, after mice received 10 shocks. Treadmill session was terminated if the mice failed to return to the treadmill after 3 consecutive attempts within the last minute of running. Aerobic capacity is expressed as total time or distance run (m) during the test. During metabolic treadmill experiments, the same protocol was followed but using Columbus Instruments metabolic treadmills to allow for measurement of gas exchange during the exercise testing.

#### Metabolic Cages

Throughout the calorimetry studies, a standard 12-hour light/dark cycle was maintained. Prior to data collection, all animals were weighed and acclimated to either control or SAAR diet for three days. Mice were placed in metabolic cages, and measurements began for seven consecutive days. Energy expenditure was determined using a computer-controlled indirect calorimetry system (PromethionH, Sable Systems, Las Vegas, NV) as published (Grobe, 2017). Animals had unlimited access to food and water throughout the study. XYZ beam arrays (BXYZ-R, Sable Systems, Las Vegas, NV) were used to record ambulatory activity and position, and respiratory gasses were measured using an integrated fuel cell oxygen analyzer, a spectro-photometric CO2 analyzer, and a capacitive water vapor partial pressure analyzer (GA3, Sable Systems, Las Vegas, NV). Oxygen consumption and CO2 production were monitored for 1-minute at 5-minute intervals. The respiratory quotient (RQ) was determined by dividing CO_2_ production by O_2_ consumption. The Weir equation was used to calculate energy expenditure: Kcal/hr = 60*(0.003941*VO_2_ + 0.001106*VCO_2_). MetaScreen v. 2.5 was used to coordinate data acquisition and instrument control, and raw data was processed using ExpeData v. 1.8.5 (Sable Systems, Las Vegas, NV) via an analysis macro that detailed all aspects of data transformation.

#### Body composition and Food Intake

Body mass was determined by daily measurement at approximately ZT22. Lean and fat mass were measured in awake mice using an EchoMRI 100H body composition analyzer.

#### RNA Extraction and Quantitative RT-PCR

RNA of tissues was extracted using a RNeasy Kit according to the manufacturer’s instructions (QIAGEN, 74034). RNA purity and concentration were assessed via a spectrophotometer (Tecan, Spark or NanoDrop, ThermoFisher). RNA was reverse-transcribed to cDNA by High Capacity cDNA Reverse Transcription Kit (Thermo Fisher, 43-688-13). A SYBR Green-based master mix (ThermoFisher Scientific, A25778) was used for real-time qPCR analysis with primers listed in Table S2. To compensate for variations in RNA input and efficiency of reverse-transcription, RPLP and Actin were used as a housekeeping gene. The delta-delta CT method was used to normalize the data.

#### RNA Sequencing and Differential Gene Expression Analysis

RNA sequencing was performed by Novogene. The quality and quantity of isolated RNA and final libraries were determined using Qubit Fluorometer and Tapestation (Agilent, Waldbronn, Germany). Sequencing libraries were prepared following SMARTerÒ Universal Low Input RNA Kit for Sequencing. Briefly, total RNA samples (0.25–10 ng) were reverse-transcribed using random priming into double-stranded cDNA in the presence of a template switch oligo (TSO). Ribosomal cDNA was cleaved by ZapR in the presence of the mammalian-specific R-Probes. Remaining fragments were enriched with a second round of PCR amplification using primers designed to match Illumina adapters. The product is a smear with an average fragment size of approximately 360 bp. The libraries were normalized to 10nM in Tris-Cl 10 mM, pH8.5 with 0.1% Tween 20. Read quality was assessed using FastQC. Alignment to the GRCm38 mouse reference genome was performed using the align function and annotation was performed using the featureCounts function from the Rsubread package (Liao et al., 2019). Genes were filtered based on minimum expression (> 5 counts per million in at least 5 samples). Differential gene expression was computed using a negative binomial model implemented in the DESeq and limma packages (Love et al., 2014; Ritchie et al., 2015). Significantly differentially expressed genes were defined as a p-value < 0.01 with a false discovery ratio (FDR) < 0.1. FDR values were calculated using the Benjamini–Hochberg method. Gene ontology enrePathway analysis was performed using the clusterProfiler R package (Wu et al., 2021). Over-representation analysis was performed using the differentially expressed genes (DEGs). Geneset-enrichment was performed using 3 databases: GO Biological process (BP), KEGG pathway and Reactome pathway. P-values were corrected for multiple testing using the Benjamini-Hochberg procedure and adjusted p-values < 0.05 were considered significant. Complex heatmaps were generated using the ComplexHeatmap package for R (Gu, 2022).

#### Immunoblot Analysis

Tissues were collected and lysed with [50 mM Tris–HCl pH 7.0, 270 mM sucrose, 5 mM EGTA, 1 mM EDTA, 1 mM sodium orthovanadate, 50 mM glycerophosphate, 5 mM sodium pyrophosphate, 50 mM sodium fluoride, 1 mM DTT, 0.1% Triton X-100 and a complete protease inhibitor tablet (Roche Applied Science)]. Lysates were centrifuged at 10000 g for 10 min at 4C. Supernatant was collected, and protein concentration was measured using the Pierce BCA protein assay kit (23225, ThermoFisher Scientific). 5-10 mg of total protein was loaded in a 15-well precast, gradient gel (456-8086, Bio-Rad). Proteins were transferred onto a PVDF membrane (Bio-rad, 170-4156) with a semi-dry or wet system and subsequently blocked for 1 h at room temperature with 5% milk in 0.1% TBS-Tween. Membranes were incubated overnight at 4C with primary antibodies listed in Key Resources Table. The appropriate HRP-linked secondary antibodies (see Key Resources Table) were used for chemiluminescent detection of proteins. Membranes were scanned with a Chemidoc imaging system (Bio-rad) and quantified using ImageJ software.

#### Immunohistochemistry and Histology

EDL or soleus muscle samples were harvested and embedded in Tissue-Tek and frozen in liquid N2-cooled isopentane and stored at -80 C until further use. Frozen muscle cross sections (7-10 μm) were made using a cryostat (Leica CM 1950) and collected on Superfrost Ultra Plus slides (Thermo Fisher Scientific). After acclimatizing to room temperature for approximately 15 min, skeletal muscle cryosections (10 µm) were fixed in 4% PFA for 10 min, washed three times with PBS and subsequently incubated for 1 h in blocking buffer (PBS with 10% donkey serum) at room temperature. Thereafter, samples were incubated overnight at 4C with primary antibodies diluted in blocking buffer with or without addition of 0.1% Triton X-100. Slides were subsequently washed in PBS and incubated for 1 h in blocking buffer with the appropriate secondary antibodies at 1:250 dilution. Nuclei were stained with Hoechst.

Images were captured with a Zeiss Axio observer Z.1 or an Olympus confocal microscope (FV1200). Fiber cross-sectional area was automatically determined on laminin-stained sections with the Muscle J plugin for ImageJ software (Mayeuf-Louchart et al., 2018). In the axitinib experiment, vascular density (% CD31+ area) was quantified within the whole tissue with ImageJ software after threshold processing on 20x images acquired with a Nikon Eclipse Ti2 microscope.

#### Stable isotope infusions in mice

For intravenous infusions, U-^13^C_18_-Linolate (Cambridge Isotope Laboratories) was prepared at a 2mM concentration in saline + 1mM BSA. Mice underwent surgery to insert a jugular vein catheter and were allowed to recover for at least one week before experiments. The infusion setup (Instech Laboratories) included a swivel and tether to allow the mouse to move around the cage freely. Infusion rate was set to 0.3 μL/min and tracer infused for 90 minutes followed by tail blood collection and tissue harvesting. Fasted infusions were collected at 5PM 8 hours after chow removal (started infusion at 2:30PM).

#### Metabolite extraction of serum

Serum (3 μl) was extracted with cold 100% methanol (40X), vortexed, and incubated on dry ice for 30 min. Then, the extract was centrifuged at 20,000 x g for 20 minutes at 4°C and supernatant was transferred to tubes containing -fold excess 100% methanol, vortexed and incubated on dry ice for 30 min. Then, the extract was centrifuged at 20,000 x g for 20 minutes at 4°C and supernatant was transferred to tubes for LC-MS analysis.

#### Metabolite extraction of tissues

Frozen tissue pieces were pulverized using a Cryomill (Retsch) at cryogenic temperature. Ground tissue was weighed (10–20 mg) and transferred into a precooled tube for extraction. Soluble metabolites extraction was done by adding −20 °C 40:40:20 methanol:acetonitrile:water to the resulting powder (40 μl solvent per mg tissue). Samples were vortexed for 10 seconds, cooled at 4°C (on wet ice) for 20 minutes and then centrifuged at 4 °C at 20,000 x *g* for 30 minutes. Supernatant was transferred to LC–MS vials for analysis.

#### Metabolite measurement by LC-MS

LC−MS analysis for soluble metabolites was achieved on a quadrupole-orbitrap mass spectrometer (Thermo Scientific): the Q Exactive PLUS hybrid, Exploris 240 or Exploris 480. Each mass spectrometer was coupled to hydrophilic interaction chromatography (HILIC) via electrospray ionization. To perform the LC separation of serum and tissue samples, an XBridge BEH Amide column (150 mm × 2.1 mm, 2.5 μM particle size, Waters) was used with a gradient of solvent A (95%:5% H2O: acetonitrile with 20 mM ammonium acetate, 20 mM ammonium hydroxide, pH 9.4), and solvent B (100% acetonitrile). The gradient was 0 minutes, 85% B; 2 minutes, 85% B; 3 minutes, 80% B; 5 minutes, 80% B; 6 minutes, 75% B; 7 minutes, 75% B; 8 minutes, 70% B; 9 minutes, 70% B; 10 minutes, 50% B; 12 minutes, 50% B; 13 minutes, 25% B; 16 minutes, 25% B; 18 minutes, 0% B; 23 minutes, 0% B; 24 minutes, 85% B; 30 minutes, 85% B. The flow rate was 150 μl min^−1^, an injection volume of 10 μl for serum samples and 5 μl for tissue samples, and column temperature was 25°C. MS full scans were in negative or positive ion mode with a resolution of 140,000 at *m/z* 200 and scan range of 70–1,000 *m/z*. The automatic gain control (AGC) target was 1 × 10^6^. LC-MS peak files were analyzed and visualized with El-MAVEN (Elucidata) using 5 ppm ion extraction window, minimum peak intensity of 1 x 10^5^ ions, and minimum signal to background blank ratio of 2. For infusion experiments, the software package Accucor was used to correct for metabolite labeling from natural isotope abundance.

#### Circulating flux measurements

To measure the circulating (whole-body) flux of a metabolite, we infused U^13^C-labeled form of the metabolite. At pseudo-steady state, we measured the mass isotope distribution of the metabolite in serum and the intact tracer circulatory flux (*F*_circ_) was calculated as previously described. The fraction of the fully labeled tracer (i.e., the infused form), L_M+C_ (for example, linolate is M+18 due to having 18 carbon atoms) was used:

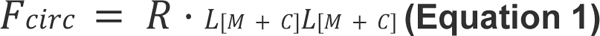

where R is the infusion rate of the labeled tracer. Since the circulatory flux is a pseudo-steady state measurement, for minimally perturbative tracer infusions, production flux is approximately equal to consumption flux of the metabolite and thus *F*_circ_ reflects both the circulating production and consumption fluxes of the infused metabolite.

#### Lipid extraction of tissues

Frozen tissue pieces were pulverized using a Cryomill (Retsch) at cryogenic temperature. Soluble lipid extraction was done by adding 50% methanol:50% H_2_O and chloroform 2:1 to the samples. Samples were incubated at 4°C (on wet ice) for 10 min and then centrifuged at 4 °C at 20,000 x *g* for 5 min. The bottom layer was extracted using glass hamilton syringes and transferred to glass vials for further processing. The first extraction step was repeated and the chloroform was evaporated using a nitrogen gas manifold. Samples were reconstituted in 1:1:1 methanol, acetonitrile, isopropyl alcohol for analysis.

#### Lipid measurement by LC-MS

Lipids were analyzed using a Vanquish Horizon UHPLC System (Thermo Scientific) coupled to a Q Exactive Plus mass spectrometer (Thermo Scientific). Agilent Poroshell 120 EC-C18 column (particle size, 2.7 μm; 150 mm (length) × 2.1 mm (i.d.)) was used for separation.

Column temperature was 25 °C. Mobile phases A = 1 mM ammonium acetate and 0.2% (v/v) acetic acid in 90:10 (v/v) water:methanol and B = 1 mM ammonium acetate and 0.2% (v/v) acetic acid in 98:2 (v/v) isopropanol:methanol were used for ESI positive mode. The linear gradient eluted from 25% B (0.0–2.0 min), 25% B to 65% B (2.0–4.0 min), 65% B to 100% B (4.0–16.0 min), 100% B (16.0–20.0 min), 100% B to 25% B (20.0–21.0 min), 25% (21.0 – 25.0 min). The flow rate was 0.15 mL/min. The sample injection volume was 5 μL. ESI source parameters were as follows: spray voltage, 3200 V or −2800 V, in positive or negative modes, respectively (arb = arbitrary units); sheath gas, 35 arb; aux gas, 10 arb; sweep gas, 0.5 arb; ion transfer tube temperature, 300 °C; vaporizer temperature, 35 °C. LC–MS data acquisition was operated under full scan positive mode for all samples. The full scan was set as: orbitrap resolution, 70,000 at m/z 200; AGC target, 3e6 arb; maximum injection time, 250 ms; scan range, 265–1150 m/z. LC-MS peak files were analyzed and visualized with El-MAVEN (Elucidata) using 5 ppm ion extraction window, minimum peak intensity of 1 x 10^5^ ions, and minimum signal to background blank ratio of 2. For infusion experiments, the software package Accucor was used to correct for metabolite labeling from natural isotope abundance.

#### Ex vivo beta oxidation

After seven-day dietary treatment mice fed either a Con or SAAR diet were sacrificed and EDL and soleus dissected, weighed and immediately put on ice in low glucose DMEM (Thermo Fisher Scientific). To start the assay, muscles are transferred to low glucose DMEM media containing 2% fatty acid free BSA, 0.25 mM carnitine and 2µCi/ml [9,10-3H]-palmitic acid (NET53100, PerkinElmer, Zaventem, Belgium). Tissues were incubated for 3 h in culture medium at 37 C and 5 % CO_2_, after which the supernatant was taken, and 10% Trichloroacetic acid (TCA) added and incubated at room temperature for 15 min. Samples were spun down at max speed for 10 min before 5% TCA was added followed by 10% BSA in TE buffer. After 15 min of incubation samples were spun down again and the supernatant was incubated with Chloroform:Methanol (2:1) and KCl:HCl was added. Samples were spun down on last time, before the supernatant was transferred into scintillation vials and 3H labeling was determined using a b-counter. CPM values were background subtracted and normalized to mg wet weight of the tissue.

#### Isolation of Endothelial Cells

Primary ECs from skeletal muscle (mECs) were isolated from adult WT and EC^CD36-/-^ littermates. Mice were euthanized, all hind-limb muscles were immediately dissected, and muscles were minced in a Petri dish on ice using a surgical blade. Next, the minced muscle tissue was enzymatically digested in digestion buffer containing 2 mg/mL Dispase II (D4693, Sigma-Aldrich), 2 mg/mL Collagenase IV (17104019, Thermo Fisher Scientific) and 2 mM CaCl_2_ in PBS at 37°C for 40 min, with gentle shaking every 10 min. The reaction was stopped by adding an equal volume of 20% FBS in HBSS and the suspension was passed through a series of 100-μm cell strainers (Corning) and 70-μm cell strainers (Corning) to remove tissue debris. After a series of centrifugation and washing steps, the heterogeneous cell population was purified by FACS.

#### Flow Cytometry

Cells were incubated in PBS with the fixable viability dye eFluor® 780 (65-0865-14, eBioscience) before antibody staining. Prior to surface staining with antibodies, Fc gamma receptors were blocked by incubating cells with anti-CD16/CD32 antibodies (2.4G2, homemade). Thereafter, cells were incubated with the appropriate primary antibodies (CD45, CD31, CD36) diluted in FACS buffer (DPBS + 2% FCS) and subsequently incubated with antibodies for 30 min on ice. For EdU proliferation experiments, cells from EdU-injected mice were first stained with antibodies for cell surface markers and subsequently labeled with the click-iT plus EdU Alexa Fluor® 488 Flow Cytometry Assay Kit (Life Technologies) according to the manufacturer’s instructions. Cells were analyzed with a LSRFortessa (BD Bioscience) flow cytometer or sorted using a FACS Aria III (BD Bioscience) sorter. Data were analyzed using FlowJo 10 software (Tree Star). A complete list of all antibodies and staining reagents used can be found in Key Resources Table. The gating strategies used for flow cytometry plots are shown in Figure S3.

#### Enzyme-Linked Immunosorbent Assay (ELISA)

Skeletal muscle tissue samples (10-15 mg) were homogenized with a tissue homogenizer (Omni THq) in ice-cold lysis buffer (1:15 w/v) as described above. Homogenates were centrifuged at 10000 g for 10 min at 4C, and VEGF was measured in the supernatants using the Mouse VEGF Quantikine ELISA Kit (R&D System, MMV00) according to the manufacturer’s protocol.

### QUANTIFICATION AND STATISTICAL ANALYSIS

The images presented in the manuscript are representative of the data (quantification of image is approximately the group average) and the image/staining quality. All data represent mean ± SD. GraphPhad Prism software (version 8.0.0) was used for statistical analyses. Investigators were always blinded to group allocation. When comparing two group means, Student’s t test was used in an unpaired two-tailed fashion. For more than two groups, one-way ANOVA with Tukey’s multiple comparisons test was used and for experimental set-ups with a second variable, two-way ANOVA with Sidak’s multiple comparisons test was used. The statistical method used for each experiment is indicated in each figure legend. No experiment-wide multiple test correction was applied. p > 0.05 is considered non-significant. p < 0.05 is considered significant.

## Supplemental Figures

**Supplemental Figure 1.**
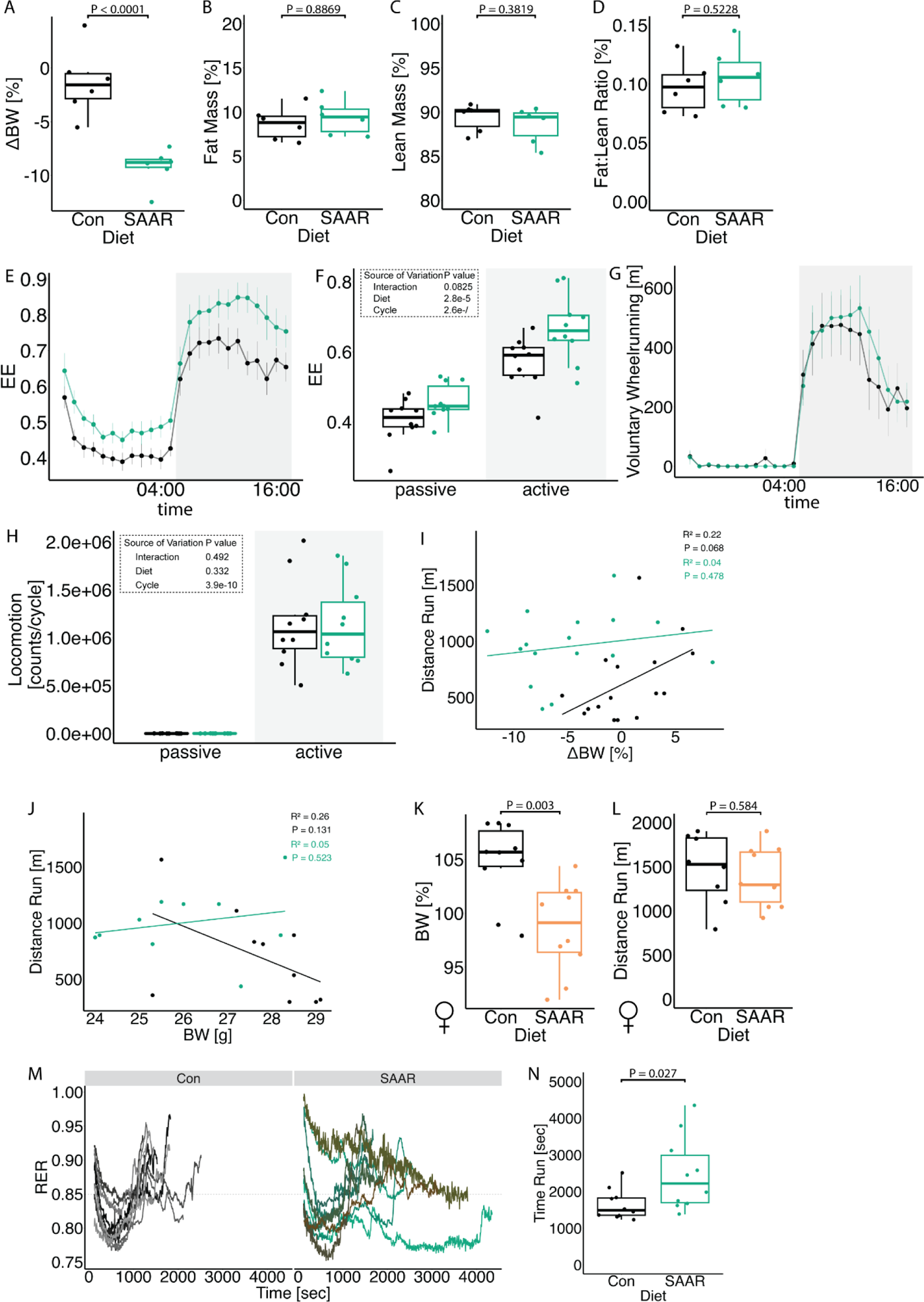
Short-term SAAR induces shifts in metabolism and increases endurance exercise capacity in young, sedentary male mice. A. Percent change in body weight (n = 6) of male mice given *ad libitum* access to sulfur amino acid restricted (SAAR) versus control (Con) diet for seven days. B. Percent fat mass of total body weight measured using ECHO MRI (n = 6) of male mice given *ad libitum* access to SAAR versus Con diet on day seven. C. Percent lean mass of total body weight measured using ECHO MRI (n = 6) of male mice given *ad libitum* access to SAAR versus Con diet on day seven. D. Fat:lean mass ratio calculated from A and B (n = 6) of male mice given *ad libitum* access to SAAR versus Con diet on day seven. E. Percent change in body weight (n = 8) of female mice given *ad libitum* access to SAAR versus Con diet after seven days on the diet. F. Distance ran during a one-time maximal endurance test (n = 8) of female mice given *ad libitum* access to SAAR versus Con diet. G. Sable systems indirect calorimetry measurements of energy expenditure (kcal, EE) over a 24 h period (n = 10) of mice given *ad libitum* access to SAAR versus Con diet for seven days and H. the average EE during a 12 h–12 h light–dark cycle (n = 10) of male mice given *ad libitum* access to SAAR versus Con diet on day seven. I. Sable systems indirect calorimetry measurements of sum of beam breaks (counts/cycle, Locomotion) over a 24 h period (n = 10) of male mice given *ad libitum* access to SAAR versus Con diet for seven days. J. Sable systems indirect calorimetry measurements of voluntary wheel running behavior over a 24 h period (n = 10) of male mice given *ad libitum* access to SAAR versus Con diet for seven days. K. Linear regression showing the relationship between relative change in body weight versus distance ran (n = 16) of male mice given *ad libitum* access to SAAR versus Con diet on day seven. R^2^ coefficient was calculated using Pearson’s method. L. Linear regression showing the relationship between absolute body weight versus distance ran (n = 16) of male mice given *ad libitum* access to SAAR versus Con diet on day seven. R^2^ coefficient was calculated using Pearson’s method. M. RER trajectory over time in seconds during a one-time maximal endurance test on metabolic treadmills (Harvard Apparatus) (n = 10) of male mice given *ad libitum* access to SAAR versus Con diet on day seven. N. Time ran in seconds during a one-time maximal endurance test measured on metabolic treadmills (n = 10) of male mice given *ad libitum* access to SAAR versus Con diet. All data is shown as mean and error bars indicate SD unless otherwise noted; p values indicate the significance of the difference by Student’s t test or two-way ANOVA with Sidak’s multiple comparisons test between diets or diet and cycle (indirect calorimetry); significance is determined by a p value of p < 0.05. For linear regressions r squared Pearson’s coefficient was calculated.

**Supplemental Figure 2.**
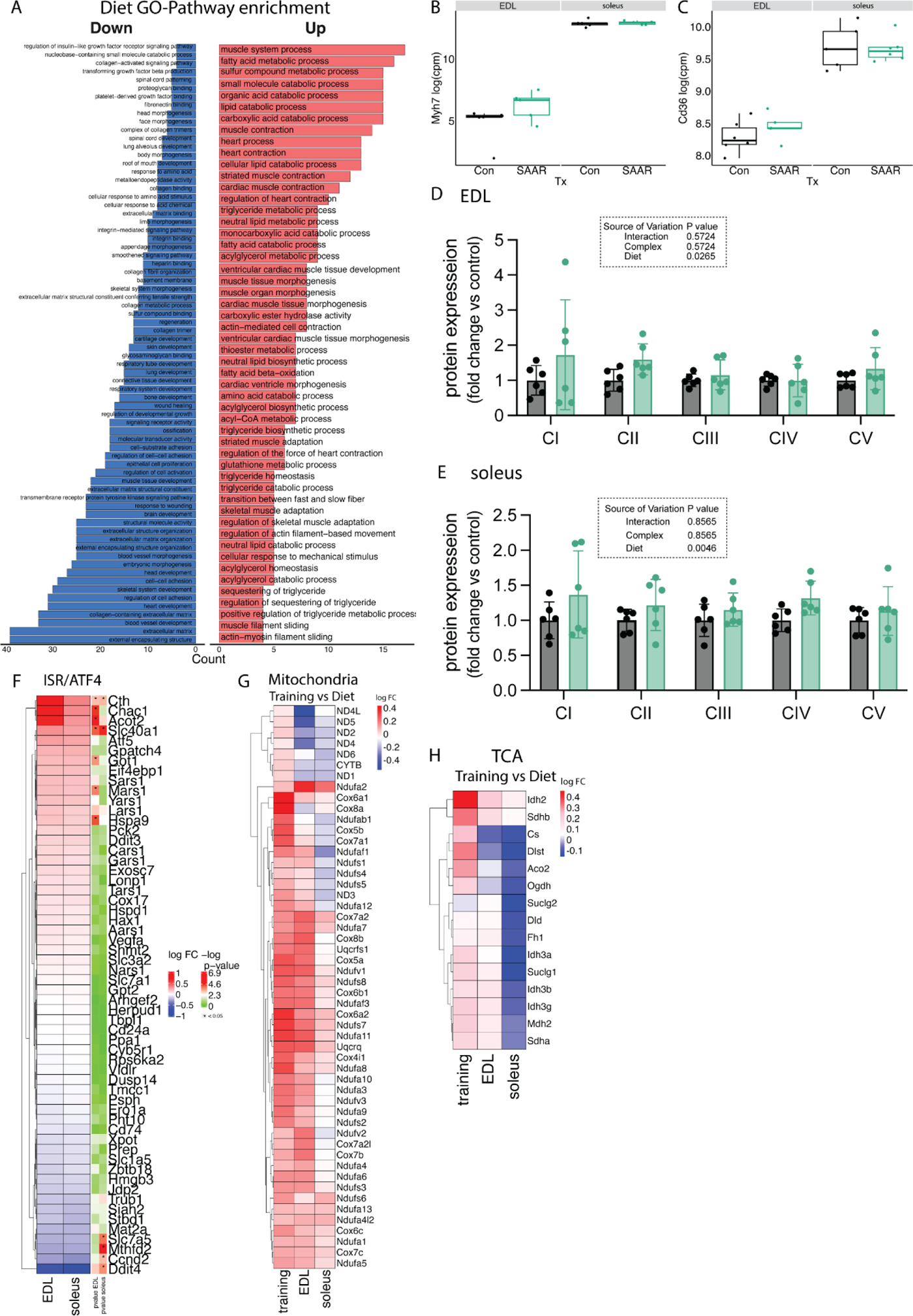
Transcriptomics across muscle depots reveal metabolic shift from glycolytic toward oxidative. A. Pathway enrichment analysis comparing main effects of diet from bulkRNA sequencing (n = 6) of male mice given *ad libitum* access to sulfur amino acid restricted (SAAR) versus control (Con) diet on day seven showing all significantly increased or decreased pathways. B. Normalized count values of Myh7 in EDL and soleus from bulkRNA sequencing (n = 6) of male mice given *ad libitum* access to SAAR versus Con diet on day seven. C. Normalized count values of Cd36 in EDL and soleus from bulkRNA sequencing (n = 6) of male mice given *ad libitum* access to SAAR versus Con diet on day seven. D. Quantification of relative protein abundance normalized to vinculin of all five complexes of the electron transport chain from blots shown in figure 2E of both EDL and soleus (n = 5). E. Fold changes of transcripts associated with known dietary SAAR and integrated stress response (ISR) target genes (Torrence et al., 2021) after SAAR when compared to Con. F. Fold changes after Training (Furrer et al., 2023) and SAAR when compared to Con of specific genes associated with mitochondrial matrix. G. Fold changes after Training (Furrer et al., 2023) and SAAR when compared to Con of specific genes associated with TCA cycle as identified in figure 2E. All data is shown as mean and error bars indicate SD unless otherwise noted; p values indicate the significance of the difference by Student’s t test or two-way ANOVA with Sidak’s multiple comparisons test between diets or diet and complexes; significance is determined by a p value of p < 0.05.

**Supplemental Figure 3.**
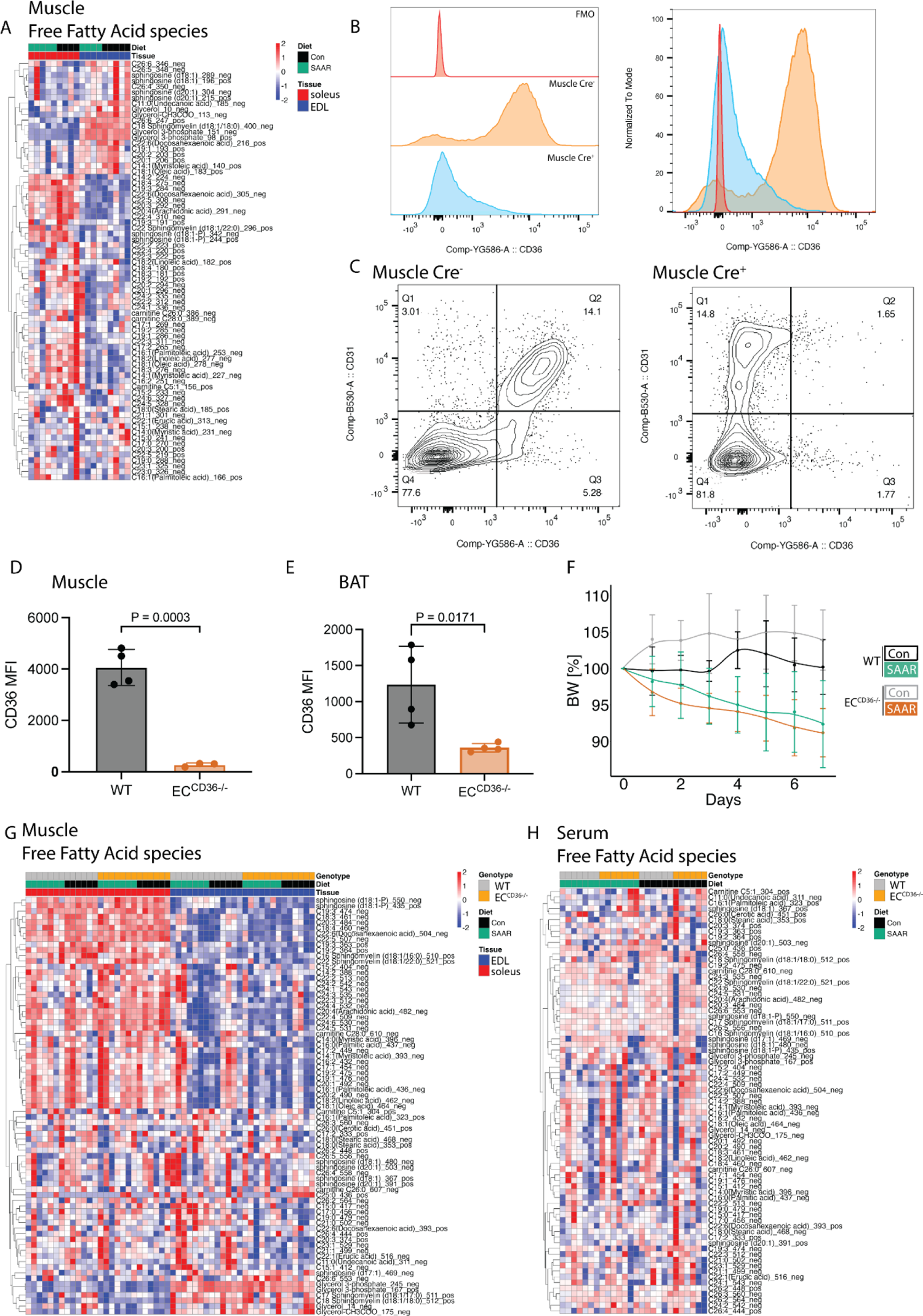
SAAR increases muscle lipid flux without altering lipid pool sizes. A. Heatmap of differentially abundant free fatty acid species in muscles (n= 3-4) of male mice given *ad libitum* access to sulfur amino acid restricted (SAAR) versus control (Con) diet for seven days. B. Representative histograms of CD36^+^ endothelial cells (EC) in the muscle and C. gating strategy for CD36 positive EC (CD45^-^, CD31^+^, CD36^+^) isolated from muscle of male WT (Cre^-^) and EC^CD36-/-^ (Cre^+^) mice. D. EC^CD36-/-^ KO efficiency was confirmed by FACS analysis of CD31^+^/CD36^+^ MFI in muscle or E. brown adipose tissue (BAT) (n = 4) of male WT or EC^CD36-/-^ mice. F. Daily body weight trajectories shown in percent of starting body weight (n = 8/group) over time of male WT and EC^CD36-/-^ mice given *ad libitum* access to SAAR versus Con diet for seven days. G. Heatmap of differentially abundant free fatty acid species in muscles of male WT or EC^CD36-/-^ mice (n = 5) fed a Con or SAAR diet for seven days. H. Heatmap of differentially abundant free fatty acid species in serum of male WT or EC^CD36-/-^ mice (n = 5) fed a Con or SAAR diet for seven days. All data is shown as mean and error bars indicate SD unless otherwise noted; p values indicate the significance of the difference by Student’s t test between diets, or two-way ANOVA with Sidak’s multiple comparisons test between diets and muscle or genotype; significance is determined by a p value of p < 0.05.

**Supplemental Figure 4.**
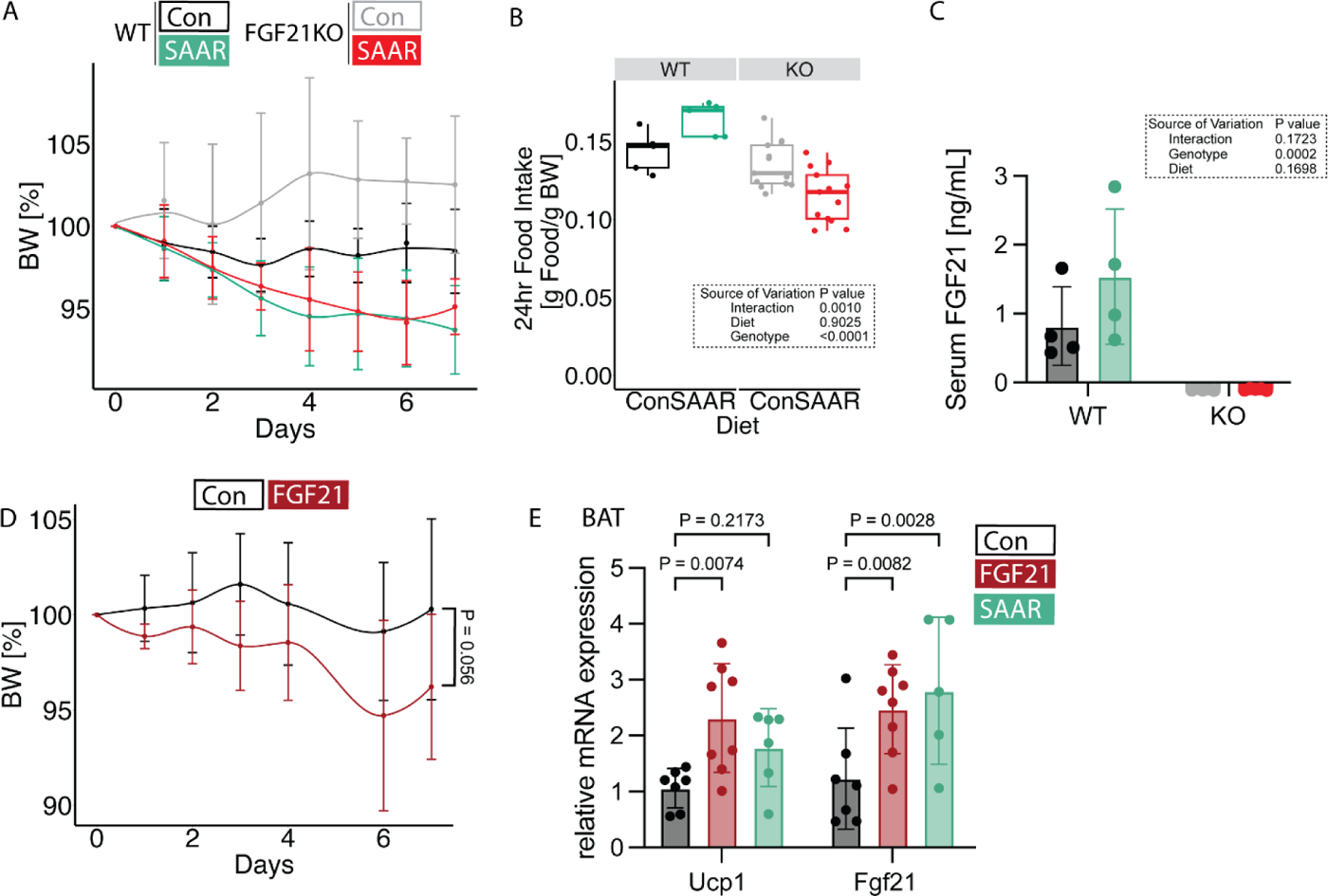
FGF21 is dispensable for running phenotype after SAAR in male mice. A. Daily body weight trajectories shown in percent of starting body weight (n = 4-10) over time, of male WT or FGF21KO mice given *ad libitum* access to sulfur amino acid restricted (SAAR) versus control (Con) diet for seven days. B. Food intake expressed as grams of food eaten per gram of mouse body weight within a 24 hr period (n = 4-10) of male WT or FGF21KO mice given *ad libitum* access to SAAR versus Con diet for seven days. C. Serum FGF21 concentrations of male WT or FGF21 KO mice given *ad libitum* access to SAAR versus Con diet for seven days determined using an ELISA. D. Daily body weight trajectories over time shown in percent when compared to starting body weight (n = 8) of NaCl or recombinant FGF21 treated male mice for seven days. E. Fgf21 and Ucp1 mRNA levels in brown adipose tissue (BAT) of male mice given *ad libitum* access to SAAR versus Con diet or mice treated with recombinant FGF21 for seven days All data is shown as mean and error bars indicate SD unless otherwise noted; p values indicate the significance of the difference by Student’s t test between diets, or two-way ANOVA with Sidak’s multiple comparisons test between diets and genotype; significance is determined by a p value of p < 0.05.

**Supplemental Figure 5.**
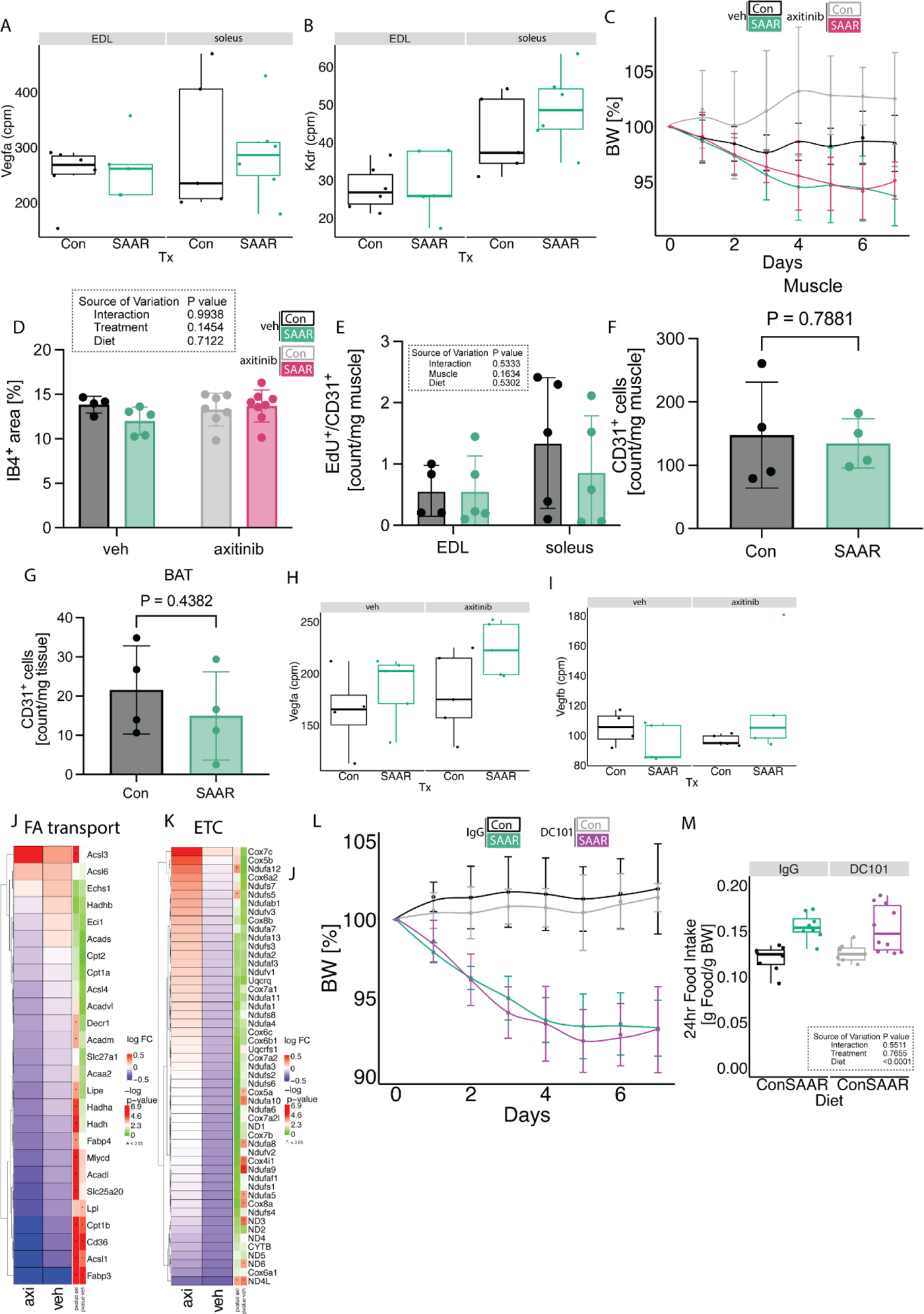
Inhibition of VEGFR signaling prevents endurance exercise phenotype without induction of angiogenesis. A. Normalized count values of Vegfa in EDL and soleus from bulkRNA sequencing (n = 6) of male mice given *ad libitum* access to sulfur amino acid restricted (SAAR) versus control (Con) diet on day seven. B. Normalized count values of Kdr in EDL and soleus from bulkRNA sequencing (n = 6) of male mice given *ad libitum* access to SAAR versus Con diet on day seven. C. Daily body weight trajectories shown in percent of starting body weight (n = 10) over time of male mice given *ad libitum* access to SAAR versus Con diet for seven days, treated with either vehicle (veh) or axtinib by oral gavage. D. Quantification of IB4^+^ area of EDL muscle of male mice fed a Con or SAAR for seven days treated with veh or axitinib (n = 5-8). E. Cell counts of EdU^+^/CD31^+^ double positive cells per mg tissue in muscle from male mice given *ad libitum* access to SAAR versus Con diet for seven days and injected with EdU to label cell proliferation, determined by flow cytometry. F. Cell counts of CD31^+^ positive cells per mg tissue in muscle or G. brown adipose tissue (BAT) or of male mice given *ad libitum* access to SAAR versus Con diet for seven days, determined by flow cytometry. H. Normalized count values of Vegfa in EDL treated with veh or axitinib from bulkRNA sequencing (n = 5) of male mice given *ad libitum* access to SAAR versus Con diet on day seven. I. Normalized count values of Kdr in EDL treated with veh or axitinib from bulkRNA sequencing (n = 5) of male mice given *ad libitum* access to SAAR versus Con diet on day seven. J. Fold changes of transcripts associated with fatty acid (FA) catabolism and transport as identified in supplementary figure 2A in both EDL treated with veh or axitinib after bulkRNA sequencing (n = 5) of male mice given *ad libitum* access to SAAR versus Con diet on day seven. K. Fold changes of transcripts associated with electron transport chain (ETC) associated genes in EDL treated with veh or axitinib after bulkRNA sequencing (n = 6) of male mice given *ad libitum* access to SAAR versus Con diet on day seven. L. Body weight trajectory over time, shown as percent of starting body weight (n = 8-10) of male mice given *ad libitum* access to SAAR versus Con diet for seven days treated with IgG or DC101 via i.p. injection every other day. M. Food intake expressed as grams of food per gram of body weight per mouse within a 24 hr period (n = 8-10) of male mice given *ad libitum* access to SAAR versus Con diet treated with IgG or DC101 via i.p. injection every other day on day seven. All data is shown as mean and error bars indicate SD unless otherwise noted; p values indicate the significance of the difference by Student’s t test between diets, or two-way ANOVA with Sidak’s multiple comparisons test between diets and muscle or treatment; significance is determined by a p value of p < 0.05.

---

Supplemental Table 1 Raw data from metabolic treadmill measurements.

Supplemental Table 2 Raw counts data from RNA sequencing analysis looking at muscle and diet interaction.

Supplemental Table 3 Raw normalized ion counts from lipidomics analysis looking at muscle and diet interaction.

Supplemental Table 4 Raw normalized ion counts from metabolomics analysis looking at muscle and diet interaction.

Supplemental Table 5 Raw normalized ion counts from metabolomics analysis looking at muscle and diet interaction in different tissues from WT and CD36^EC-/-^ mice

Supplemental Table 6 Raw counts data from RNA sequencing analysis looking at diet and axitinib treatment interaction in EDL muscle.

**Supplemental Table 7:** macronutrient composition of the control and SAAR diets used.

| Product # | A17101101 | A17101102 | A17101103 |
| --- | --- | --- | --- |
|  | Control | Mid Methionine | Low Methionine |
| **Ingredient** |  |  |  |
| Casein, Lactic | 0 | 0 | 0 |
| L-Cystine | 0 | 0 | 0 |
| L-Isoleucine | 7.6 | 7.6 | 7.6 |
| L-Leucine | 15.8 | 15.8 | 15.8 |
| L-Lysine | 13.2 | 13.2 | 13.2 |
| L-Methionine | 4.5 | 1.8 | 1.2 |
| L-Phenylalanine | 8.4 | 8.4 | 8.4 |
| L-Threonine | 7.2 | 7.2 | 7.2 |
| L-Tryptophan | 2.1 | 2.1 | 2.1 |
| L-Valine | 9.3 | 9.3 | 9.3 |
| L-Histidine-HCl-H<sub>2</sub>O | 4.6 | 4.6 | 4.6 |
| L-Alanine | 5.1 | 5.1 | 5.1 |
| L-Arginine | 6 | 6 | 6 |
| L-Aspartic Acid | 12.1 | 12.1 | 12.1 |
| L-Glutamic Acid | 38.2 | 38.2 | 38.2 |
| Glycine | 3 | 3 | 3 |
| L-Proline | 17.8 | 17.8 | 17.8 |
| L-Serine | 10 | 10 | 10 |
| L-Tyrosine | 9.2 | 9.2 | 9.2 |
| **Total L-Amino Acids** | **174.1** | **171.4** | **170.8** |
| Corn Starch | 506.2 | 506.2 | 506.2 |
| Maltodextrin 10 | 125 | 125 | 125 |
| Sucrose | 73.6 | 76.3 | 76.9 |
| Cellulose, BW200 | 50 | 50 | 50 |
| Soybean Oil | 25 | 25 | 25 |
| Lard | 20 | 20 | 20 |
| Mineral Mix S10026 | 10 | 10 | 10 |
| DiCalcium Phosphate | 13 | 13 | 13 |
| Calcium Carbonate | 5.5 | 5.5 | 5.5 |
| Potassium Citrate, 1 H<sub>2</sub>O | 16.5 | 16.5 | 16.5 |
| Sodium Bicarbonate | 7.5 | 7.5 | 7.5 |
| Vitamin Mix V10001 | 10 | 10 | 10 |
| Choline Bitartrate | 2 | 2 | 2 |
| FD&C Blue Dye #1 | 0 | 0.05 | 0 |
| FD&C Yellow Dye #5 | 0.05 | 0 | 0 |
| FD&C Red Dye #40 | 0 | 0 | 0.05 |
| **Total** | **1038.450** | **1038.450** | **1038.450** |
| gm |  |  |  |
| Protein | 174.1 | 171.4 | 170.8 |
| Carbohydrate | 714.8 | 717.5 | 718.1 |
| Fat | 45.0 | 45.0 | 45.0 |
| Fiber | 50.0 | 50.0 | 50.0 |
| gm% |  |  |  |
| Protein | 16.8 | 16.5 | 16.4 |
| Carbohydrate | 68.8 | 69.1 | 69.2 |
| Fat | 4.3 | 4.3 | 4.3 |
| Fiber | 4.8 | 4.8 | 4.8 |
| kcal |  |  |  |
| Protein | 696 | 686 | 683 |
| Carbohydrate | 2859 | 2870 | 2872 |
| Fat | 405 | 405 | 405 |
| **Total** | **3961** | **3961** | **3961** |
| kcal% |  |  |  |
| Protein | 18 | 17 | 17 |
| Carbohydrate | 72 | 72 | 73 |
| Fat | 10 | 10 | 10 |
| **Total** | **100** | **100** | **100** |
| kcal / gm | 3.8 | 3.8 | 3.8 |

A17101101-03.for

